# TRIM21 and PHLDA3 Negatively Regulate the Cross-Talk between the PI3K/AKT Pathway and PPP Metabolism

**DOI:** 10.1101/750976

**Authors:** Jie Cheng, Yan Huang, Xiaohui Zhang, Yue Yu, Wanru Zhang, Shumin Wu, Jing Jiao, Mei Wang, Linh Tran, Liuzhen Zhang, Mengyao Wang, Wenyu Yan, Yilin Wu, Fangtao Chi, Peng Jiang, Xinxiang Zhang, Hong Wu

**Author notes:** These authors contributed equally to this work (Yan Huang, Xiaohui Zhang). Corresponding author: Hong Wu: The MOE Key Laboratory of Cell Proliferation and Differentiation, School of Life Sciences, Peking-Tsinghua Center for Life Sciences and Beijing Advanced Innovation Center for Genomics, Peking University, Beijing, 100871, China.

## Abstract

PI3K/AKT signaling is known to regulate cancer metabolism but whether metabolic pathway feedbacks and regulates the PI3K/AKT pathway is unclear. Here, we demonstrate the important reciprocal cross-talks between the PI3K/AKT signal and PPP branching metabolic pathways. PI3K/AKT activation stabilizes G6PD, the rate-limiting enzyme of PPP, by inhibiting a newly identified E3 ligase TIRM21, and promotes PPP. PPP metabolites, in turn, reinforce AKT activation and further promote cancer metabolic reprogramming by blocking the expression of an AKT inhibitor PHLDA3. Knockout TRIM21 or PHLDA3 promotes the cross-talks and cell proliferation. Importantly, *PTEN* null human cancer cells and *in vivo* murine models are sensitive to anti-PPP treatments, suggesting the importance of PPP in maintaining AKT activation even in the presence of a constitutively activated PI3K pathway. Our study suggests that blockade of these reciprocal cross-talks may have a therapeutic benefit for cancers with PTEN loss or PI3K/AKT activation.

## INTRODUCTION

Metabolic reprogramming is one of the hallmarks of human cancers ^1^. Unlike normal cells, cancer cells catabolize large amounts of glucose for lactate production, even when oxygen is abundant, through a phenomenon called aerobic glycolysis or the “Warburg effect”^2^. Cancer cells express pyruvate kinase 2 (PKM2), a growth signal-sensitive version of the rate-limiting enzyme in the last step of glycolysis^3^, leading to the accumulation of glycolytic intermediates. The accumulated glycolytic intermediates can branch into pentose phosphate pathway (PPP) for nucleotide biosynthesis and NADPH production, important for cell proliferation and redox state maintenance^4^. PPP is highly upregulated in cancers to meet the needs for uncontrolled cell proliferation^5, 6^. Cancer cells are also more sensitive to PPP inhibition^7^ and chemotherapies that inhibit nucleotide biosynthesis^8^. However, the mechanisms that orchestrate glycolysis and PPP in normal and cancer cells are largely unknown.

The PI3K/AKT pathway is one of the most commonly altered signaling pathways in human cancers. The PI3K/AKT pathway converges signals from extracellular stimuli, such as growth factors, nutrients and extracellular matrix, to intracellular signaling networks that control cell growth, survival and metabolism^9–11^. PI3K/AKT promotes the Warburg effect by increasing the expression and membrane translocation of glucose transporters and the activities of enzymes involved in glycolysis. PTEN is the major tumor suppressor that antagonizes the PI3K/AKT pathway via its lipid phosphatase activity^12–14^ and therefore exerts an “anti-Warburg” effect. Indeed, overexpression of the murine *Pten* gene in a transgenic model decreases glycolysis and increases respiration^15^. However, since PTEN possesses both lipid and protein phosphatase activities as well as phosphatase-independent activities^14^, it is not clear whether the metabolic phenotype observed in the *Pten* overexpression model is solely due to its lipid phosphatase or anti-PI3K/AKT activity. It is also not clear whether PTEN loss or PI3K/AKT activation controls the PPP branching pathway in cancer metabolic reprogramming.

To answer these questions, we genetically knocked-in two cancer-associated PTEN point mutations into the endogenous *Pten* gene in embryonic stem cells (mES): the C124S mutation that is phosphatase-dead and G129E mutation that is lipid phosphatase-dead and protein phosphatase-active. These two mutant lines, together with the parental WT and *Pten* null lines^16^, allowed us to genetically separate the lipid and protein phosphatase activities as well as phosphatase-independent activity of PTEN without perturbing its level (Figure S1A and S1C). Using this true isogenic system, we conducted metabolic chasing analyses on these four cell lines and in ES cell system that mimics cancer metabolism^17, 18^.

Here, we report a novel reciprocal cross-talk between the PI3K/AKT and the PPP pathways in *Pten* mutant mES cells, which can be further confirmed in human cancer cells and *in vivo* cancer models with PTEN loss. PTEN loss or PI3K/AKT activation promotes a shift of glycolytic intermediates to the PPP branching pathway by stabilizing the rate limiting enzyme G6PD. PPP metabolites, in turn, provide positive feedback and reinforce PI3K/AKT activation via negative regulation of an AKT inhibitor PHLDA3. The positive feedback between metabolic pathway and cell signaling may have important therapeutic implications for cancers with PTEN loss and PI3K/AKT activation.

## RESULTS

### PTEN loss or PI3K activation promotes the Warburg effect by decoupling glycolysis and the TCA cycle

To fully explore the roles of PTEN in regulating cell metabolism, we measured glucose consumption in the isogenic WT, Null, CS and GE mES cells under standard ES culture condition and found that all three *Pten* mutant lines consumed more glucose and expressed higher levels of GLUT1 (Figure 1A, upper and lower left panels). All *Pten* mutant lines also secreted more lactate and had higher ECAR rates (Figure 1A, lower right panel; Figure S1B). Since all three *Pten* mutant lines lack lipid phosphatase activity (Figure S1A and S1C), this result suggests that PTEN regulates the “Warburg effect” by antagonizing the PI3K activity.

**Figure 1.**
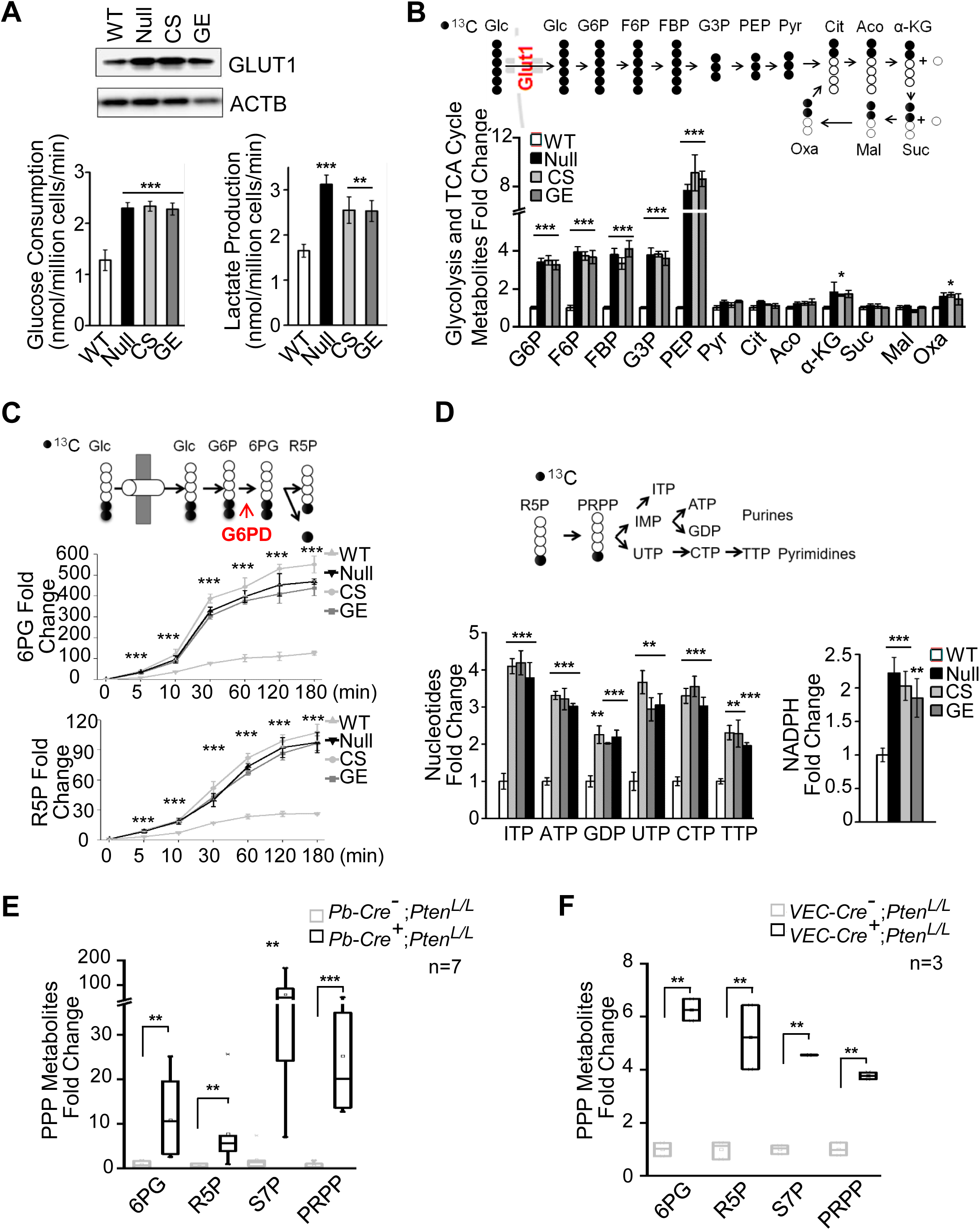
PTEN loss or PI3K activation promotes glycolysis and PPP. (A) Loss of PTEN lipid phosphatase activity increases GLUT1 levels (upper panel), glucose consumption (lower left panel) and lactate production (lower right panel) in *Pten* null, CS, and GE mES cells compared to isogenic WT cells. (B) Upper panel, a schematic illustrating [U-^13^C] glucose metabolism through glycolysis and the TCA cycle; lower panel, loss of PTEN lipid phosphatase activity increases the levels of ^13^C-labeled glycolytic intermediates from G6P to PEP in *Pten* null, CS, and GE mES cells compared to isogenic WT cells. (C) Upper panel, a schematic illustrating [1, 2-^13^C] glucose tracing into the oxidative arm of the PPP; lower panel, faster and higher levels of labeled 6PG and R5P in *Pten* null, CS, and GE mES cells compared to WT cells. (D) Upper panel, a schematic illustrating [1, 2-^13^C] glucose tracing into the nucleotide biosynthesis pathway; lower panel, increased levels of labeled nucleotides and NADPH production in *Pten* null, CS, and GE mES cells compared to WT cells. (E-F) Increased PPP metabolites in *Pten* null prostate cancer and T-ALL mouse models. Metabolites in PPP were extracted from the anterior lobes of *Pb-Cre^−^;Pten^L/L^* and *Pb-Cre^+^;Pten^L/L^* mice (E) or thymus of the *VEC^−^Cre^−^;Pten^L/L^*and *VEC^−^Cre^+^;Pten^L/L^* mice (F), then detected by LC-MS and normalized by protein concentration. Cell extracts were prepared and analyzed using LC-MS. Data are presented as fold changes, means ± SD, compared to WT cells. Each experiment was performed at least three independent times. * p< 0.05, ** p< 0.001, *** p<0.001, based on the Student’s *t*-test. See also Figure S1.

We then investigated the fate of uniformly labeled [U-^13^C]-glucose by measuring the metabolic intermediates within glycolysis and the TCA cycle (Figure 1B, upper panel)^19^. Although no obvious differences were observed in TCA metabolites among WT and *Pten* mutant lines, the levels of glycolytic intermediates from G6P to PEP were significantly increased in all *Pten* mutant lines (Figure 1B, lower panel). In contrast, pyruvate (pyr), the downstream metabolite of PEP whose production is catalyzed by the rate limiting enzyme PKM2 before TCA cycle^8, 20^, showed no significant changes (Figure 1B, lower panel). PKM2 Y105 phosphorylation is known to inhibit PKM2 activity and promote the Warburg effect and tumor growth^3, 21^. We observed significantly increased PKM2 Y105 phosphorylation and decreased PKM activity in all *Pten* mutant lines (Figure S1C), which may explain PEP accumulation and slower conversion from PEP to pyruvate. Therefore, loss of the PTEN lipid phosphatase activity or activation of the PI3K/AKT pathway controls the Warburg effect by inhibiting PKM2 activity and decoupling glycolysis and the TCA cycle.

### PI3K activation diverts glycolytic intermediates to PPP branching pathways for biosynthesis and NADPH production

Since the loss of PTEN lipid phosphatase activity led to 4-8-fold increases in the levels of glycolytic intermediates from G6P to PEP (Figure 1B), we next investigated whether these accumulated metabolites would flow to the branching metabolic pathways. We traced [1, 2-^13^C]-glucose for indicated times and quantified the amount of labeled glucose entering the oxidative arm of the PPP^19^. As shown in Figure 1C, [1, 2-^13^C]-glucose incorporated into 6PG and R5P much faster and at higher levels in *Pten* mutant lines compared to the WT cells. Furthermore, intermediates from both oxidative and nonoxidative arms (traced with [U-^13^C]-glucose), PPP-associated nucleotides and NADPH production were all significantly increased in the *Pten* mutant lines compared to the WT cells (Figure 1D; Figure S1D).

To determine whether loss of PTEN also diverts glycolytic intermediates to PPP *in vivo*, we measured PPP metabolic intermediates in the *Pten* null prostate cancer and T-ALL mouse models as they closely mimic the clinical features of these human cancers with high frequencies of PTEN mutation and PI3K pathway activation^22–25^. Consistent with the results from isogenic ES cell lines, the PPP metabolites were significantly higher in the *Pten* null cancer models as compared to their *Cre^−^* WT littermates (Figure 1E and 1F). Together, PTEN loss or PI3K activation drives the Warburg effect by reducing PKM2 activity and diverging accumulated glycolytic intermediates to the PPP branching metabolic pathways to support the needs for rapid proliferation of the *Pten* mutant cells.

### PI3K/AKT activation promotes PPP by stabilizing the rate-limiting enzyme G6PD

The underlying mechanisms of PI3K-controlled glucose transport and glycolysis have been well studied. However, little is known about how PI3K controls the PPP branching metabolic pathway^9, 26, 27^. G6PD is the rate-limiting enzyme in the PPP. Although we did not find significant differences in *G6pd* mRNA levels (Figure S2A), we did detect approximately 2- to 4-fold increases in G6PD protein levels and enzymatic activities in all *Pten* mutant mES lines (Figure 2A), suggesting that the PI3K pathway activity is responsible for regulating the G6PD protein level and activity. Indeed, treating *Pten* mutant mES cells with the PI3K inhibitor PKI-587 but not MEK inhibitor PD0325901 significantly inhibited G6PD enzyme activity in a time-dependent manner, accompanied by reductions of G6PD protein levels and PPP metabolites (Supplement Table 1) (Figure 2B; Figure 2C; Figure S2B). The potency of PKI-587 was comparable to 6-AN, an inhibitor of G6PD (Figure 2C).

**Figure 2.**
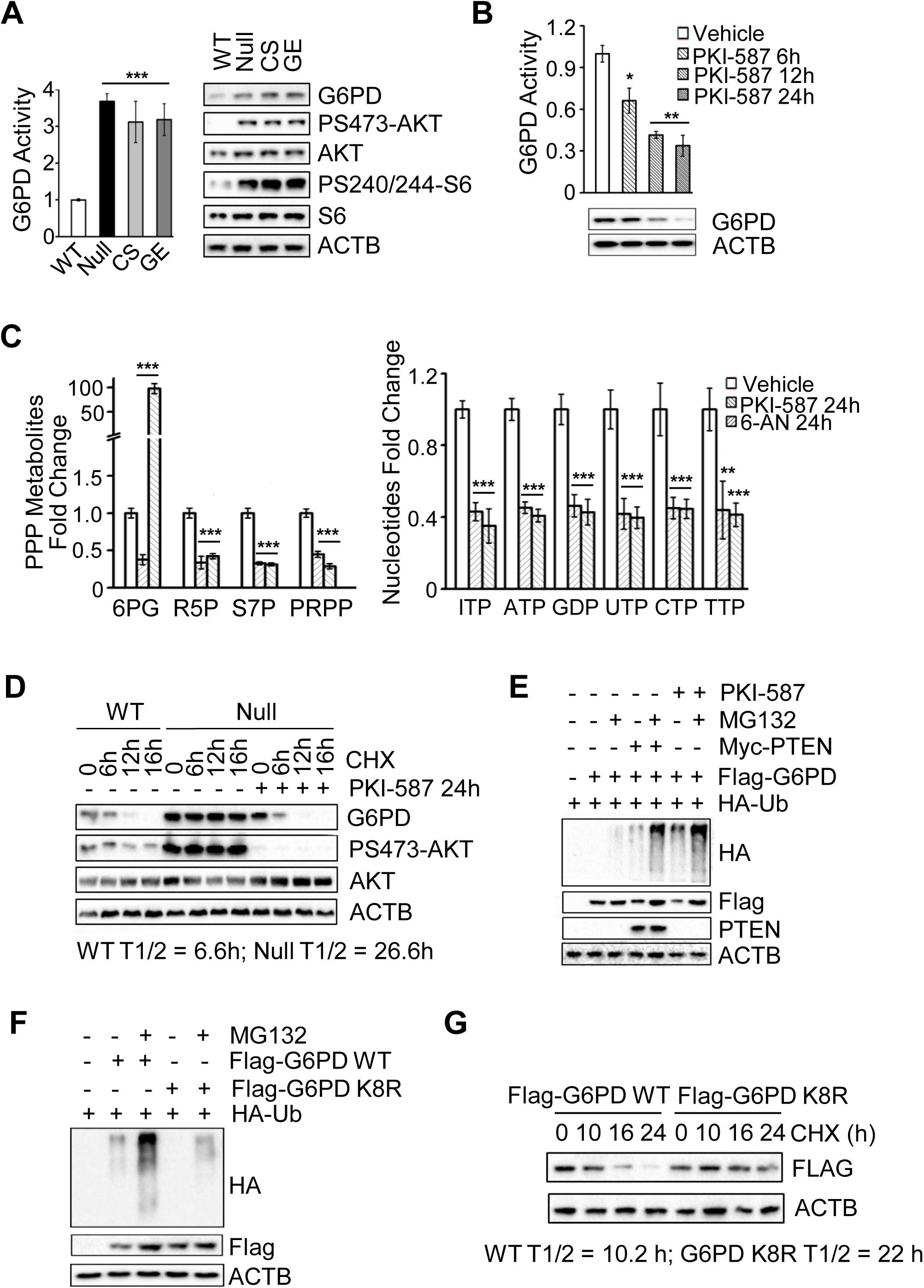
PI3K/AKT pathway promotes PPP by stabilizing the rate-limiting enzyme G6PD. (A) The enzymatic activities and protein levels of G6PD are increased in *Pten* null, CS and GE mES cells compared to WT cells. (B) Time-dependent decreases in G6PD enzymatic activity and protein levels after PKI-587 (1 µM) treatment. (C) The PPP metabolites and nucleotide productions are decreased by the PKI-587 or 6-AN treatment. *Pten* null mES cells were switching to medium containing [U-^13^C] glucose continuously cultured for 12 hours, with or without treatment of PKI-587 (1 µM) or 6-AN (100 nM) for the indicated time periods,. (D) The half-lives of G6PD in *Pten* WT and null mES cells were measured after PKI-587 (1 µM) or vehicle treatment for 24 h or treated with CHX (100 µg/ml) for the indicated time periods. Cell lysates were subjected to Western blot analysis using the indicated antibodies. (E) HEK293T cells were co-transfected with *HA-ubiquitin*, *3×Flag-G6PD* and *myc-PTEN* expression plasmids. Twenty-four hours later, cells were incubated with or without MG132 (5 µM) or PKI-587 (1 µM) for 16 h. Total cell lysates were harvested, immunoprecipitated with a Flag antibody and immunoblotted with an HA antibody and followed by Western blot analysis using the indicated antibodies. (F) HEK293T cells were co-transfected with *HA-ubiquitin*, *Flag-G6PD WT* or *Flag-G6PD K8R* expression plasmids, Twenty-four hours later, cells were incubated with or without MG132 (5 µM) for 16 h. Total cell lysates were harvested, immunoprecipitated with a Flag antibody and immunoblotted with an HA antibody and followed by Western blot analysis using the indicated antibodies. (G) The half-lives of exogenous WT and K8R G6PD were measured after CHX (100 µg/ml) treatment for the indicated time periods. Cell lysates were subjected to Western blot analysis using the indicated antibodies. Data are presented as the means ± SD and compared to WT or untreated cells. Each experiment was performed at least three independent times. * p < 0.05, ** p < 0.001, and *** p<0.001, based on Student’s *t*-test. See also Figure S2 and Supplement Table 1.

Since PI3K/AKT regulates G6PD protein instead of its mRNA, we measured the half-life of the G6PD protein in the WT and *Pten* null mES cells. PTEN loss significantly prolonged the half-life of the G6PD protein from 6.6 hours to 26.6 hours, while PKI-587 treatment completely inhibited this effect (Figure 2D). To test whether PI3K/AKT affects the stability of G6PD through the ubiquitin-mediated degradation pathway, we expressed *Myc-PTEN*, *Flag-G6PD* and *HA-ubiquitin* in 293T cells and treated the cells without or with MG132. Overexpression of PTEN or inhibition of PI3K activity significantly increased the levels of ubiquitinated G6PD, and decreased the levels of the G6PD protein that could be reverted by MG132 treatment (Figure 2E). To identify the specific residues responsible for G6PD ubiquitylation, we conducted mass spectrometry analysis and found 8 potential ubiquitylated lysines (Figure S2C). Mutating all 8 lysine (K) to arginine (R) significantly decreased ubiquitination of K8R G6PD and prolonged its half-life (Figure 2F and 2G), as compared to WT G6PD. Together, these results suggest that PTEN loss or PI3K activation promotes the PPP branching pathway by inhibiting ubiquitin-mediated degradation of G6PD.

### TRIM21 is the E3 ligase responsible for PI3K/AKT-regulated G6PD degradation

To identify the E3 ligase responsible for the G6PD degradation, we expressed *Flag-G6PD* in *PTEN* null prostate cancer PC3 cells and performed a mass spectrometry analysis of G6PD-associated proteins. Among the seven potential E3 ligases identified, TRIM21 received the highest score (Figure 3A). Reciprocal co-immunoprecipitation of endogenous proteins confirmed the physical interaction between G6PD and TRIM21 *in vivo* (Figure 3B). To test whether TRIM21 participates in G6PD ubiquitylation, we expressed *GST-TRIM21*, *Flag-G6PD* and *HA-ubiquitin* in 293T cells followed by treatment without or with MG132. As shown in Figure 3C, *TRIM21* overexpression increased G6PD ubiquitination, accompanied by decreased levels of the G6PD protein that could be reverted by MG132.

**Figure 3.**
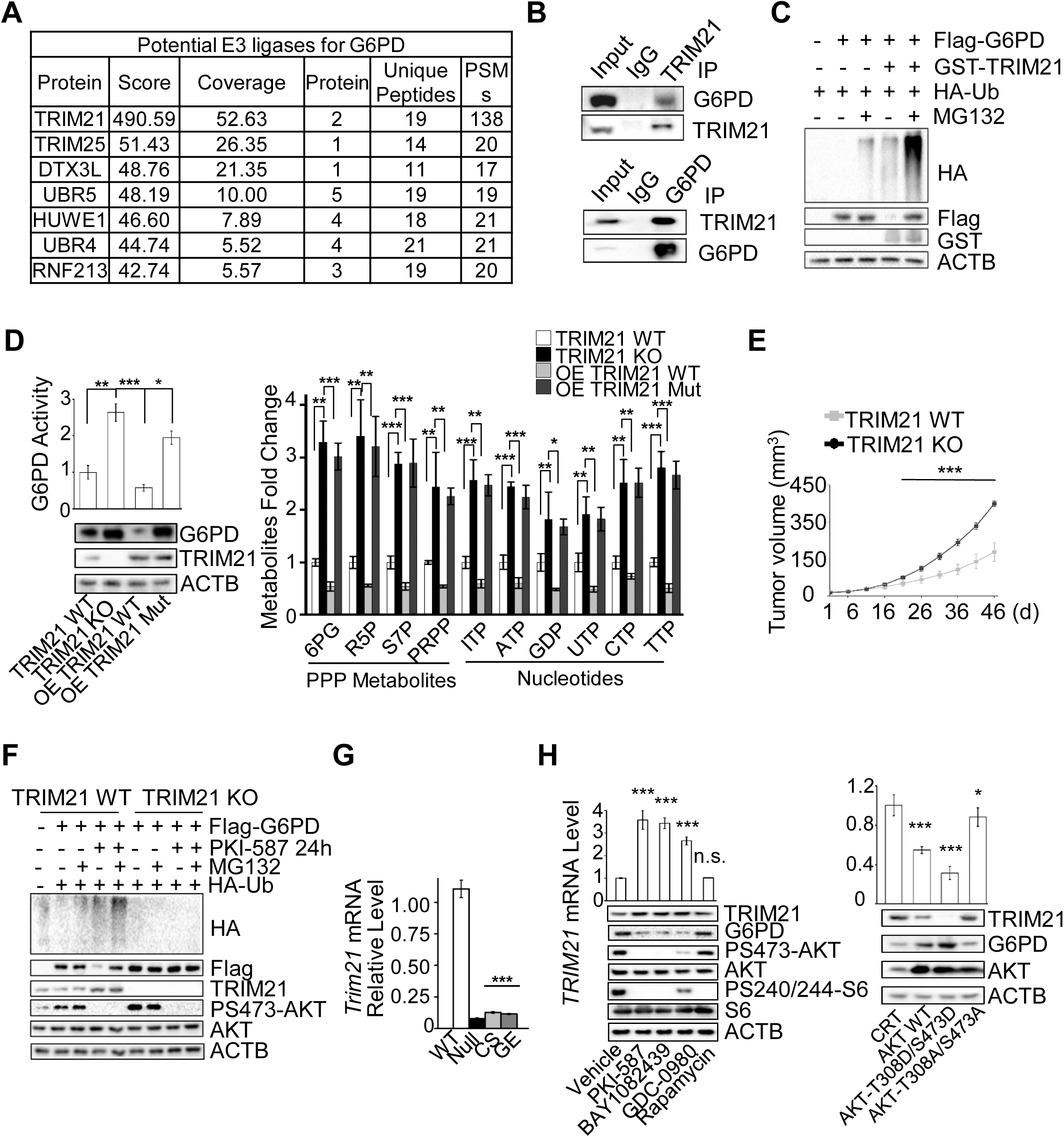
TRIM21 is the E3 ligase responsible for PI3K/AKT-regulated G6PD stability. (A) The potential G6PD E3 ligases identified by affinity MS. (B) TRIM21 physically interacts with endogenous G6PD. Cell lysates were reciprocally immunoprecipitated with either an anti-TRIM21 or -G6PD antibody, followed by immunoblotting with an anti-G6PD or -TRIM21 antibody, respectively. IgG was used as a control. (C) TRIM21 controls G6PD ubiquitination. HEK293T cells were co-transfected with *HA-ubiquitin*, *Flag-G6PD* and *GST-TRIM21* expression plasmids. Twenty-four hours later, cells were incubated with or without MG132 (5 µM) for 16 h. Total cell lysates were immunoprecipitated with a Flag antibody and immunoblotted with the indicated antibodies. (D) The enzymatic activity and protein levels of G6PD (left panel), and PPP metabolites and nucleotides (right panel) in *TRIM21* WT, knockout cells, and knockout cells re-expressing WT or E3 ligase dead mutant (C16A, C31A and H33W) *TRIM21* plasmid. (E) The growth kinetics of *TRIM21* WT- and knockout-derived xenograft *in vivo*. Equal numbers of *TRIM21* WT and knockout cells were implanted on to the bilateral flanks of nude mice. Relative tumor volumes are presented with growth kinetics. (F) TRIM21 controls G6PD ubiquitination. *TRIM21* WT and knockout cells were co-transfected with *HA-ubiquitin*, *3×Flag-G6PD* expression plasmids. Twenty-four hours later, cells were incubated with or without MG132 (5 µM) or PKI-587 (1 µM) for 16 h. Total cell lysates were harvested, immunoprecipitated with a Flag antibody and immunoblotted with an HA antibody and followed by Western blot analysis using the indicated antibodies. (G) *Trim21* mRNA levels are lower in Null, CS and GE mES cells compared to WT cells. (H) Left, PI3K/AKT inhibitors, but not rapamycin, control TRIM21 mRNA and protein levels as well as G6PD protein levels. PC3 cells were treated with PKI-587 (1 µM), BAY1082439 (5 µM), GDC-0980 (40 µM), GDC-0068 (10 µM) or rapamycin (100 nM) for 24 hours. Cell lysates were subjected to RT-PCR (upper panel) or immunoblotting with the indicated antibodies. Right, HEK293T cells were transfected with AKT-WT, AKT-T308D/S473D or AKT-T308A/S473A plasmids. Twenty-four hours later, cell lysates were subjected to RT-PCR or immunoblotting with the indicated antibodies. For metabolite measurement, cell extracts were prepared and analyzed using LC-MS. Data are presented as fold changes, means ± SD, and compared to WT cells. Each experiment was performed at least three independent times. * p < 0.05, ** p < 0.001, and *** p<0.001, based on Student’s *t*-test. See also Figure S3.

To further confirm the functional role of TRIM21 in regulating G6PD protein level, we knocked out *TRIM21.* As predicted for an E3 ligase, *TRIM21* knockout could increase both the protein and enzymatic activity of G6PD, as well as the levels of PPP metabolites (Figure 3D). Overexpression of WT *TRIM21,* but not E3 ligase dead mutant (C16A, C31A and H33W)^28^, in *TRIM21*-knockout cells could significantly reduce the levels of G6PD as well as PPP metabolites (Figure 3D), strongly supporting the notion that TRIM21 is the E3 ligase that controls G6PD degradation. Increased PPP activity in *TRIM21*-knockout cells also led increased cell growth and tumor volume *in vitro* and *in vivo*, respectively, demonstrating the significant role of TRIM21 in negatively regulating PPP and cell growth (Figure 3E; S3A and S3B).

G6PD protein is very stable in *TRIM21*-knockout cells and its ubiquitination and half-life are not altered by PKI-587 treatment (Figure 3F and S3C), suggesting that TRIM21 may be the target of PI3K/AKT signaling. Consistent with this notion, we found that *Pten* mutant mES cells had much lower *Trim21* mRNA levels than WT cells (Figure 3G). Treating *Pten* null mES cells with anti-PI3K and AKT inhibitors, but not the mTOR inhibitor rapamycin, could promote *Trim21* expression and increase TRIM21 protein levels while reduce G6PD levels (Figure 3H, left panel). Overexpression of WT AKT, especially its constitutively activated form (AKT-T308D/S473D) but not its inactivated form (AKT-T308A/S473A), could inhibit *TRIM21* transcription and decrease its protein levels, accompanied by increased G6PD protein levels (Figure 3H, right penal). These results suggest that the AKT activity is responsible for suppressing *Trim21* expression and increasing G6PD protein level and activity in *Pten* mutant ES cells.

### PI3K/AKT regulates the transcription of TRIM21 and promote PPP *in vivo* and in human cancers

To explore the roles of PI3K/AKT in regulating *Trim21* expression and PPP *in vivo*, we utilized *Pten* null prostate cancer and T-ALL mouse models. Similar to our findings in isogenic mES cell lines, we observed significant higher levels of total G6PD protein and lower levels of *Trim21* mRNA in these *Pten* null *in vivo* cancer models, as compared to their WT littermates (Figure 4A-C). Importantly, PKI-587 treatment could reduce G6PD protein levels, accompanied by increased *Trim21* expression and decreased PPP metabolites (Figure 4A-D; Figure S4A), providing strong support for this novel PI3K-regulated mechanism.

**Figure 4.**
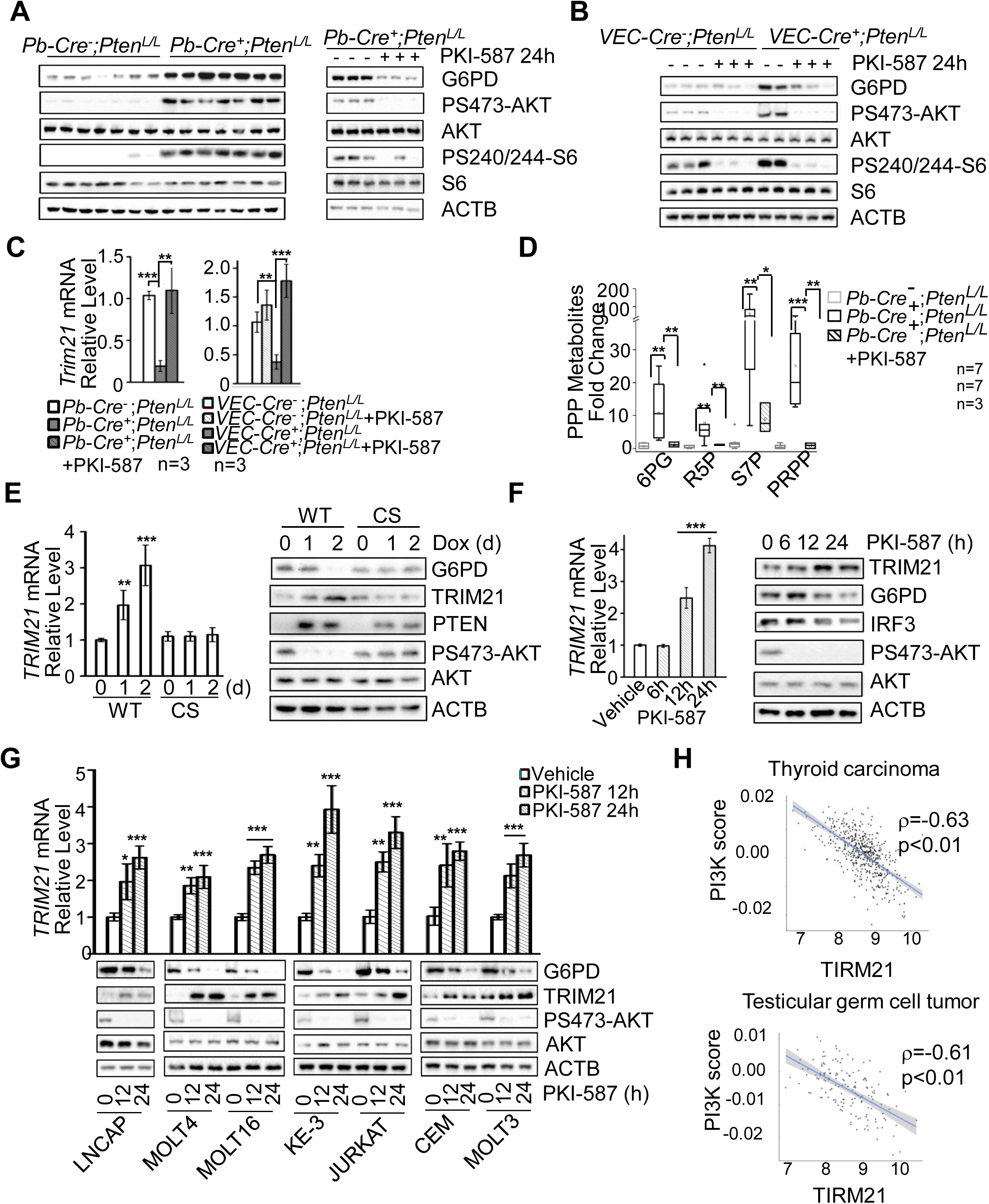
PI3K/AKT regulates TRIM21 and G6PD *in vivo* and in human cancers. (A - D) *Pten* null prostate cancer and T-ALL mouse models have higher levels of G6PD protein and PPP metabolites as well as lower levels of Trim21, which can be reverted by PKI-587 treatment. The cell lysates from the anterior lobes of *Pb-Cre^−^;Pten^L/L^* and *Pb-Cre^+^;Pten^L/L^*prostates (A) and thymic cells of *VEC^−^Cre^−^;Pten^L/L^* and *VEC^−^Cre^+^;Pten^L/L^* mice (B) without and with PKI-587 (25 mg/kg) treatment were subjected to immunoblotting with the indicated antibodies or LC-MS analysis (D) 24 h later. *Trim21* mRNA levels were measured without or with PKI-587 treatment (C). (E) *PTEN* WT/CS inducible PC3 cells were treated with Dox (2 μg/ml) for the indicated time periods. *TRIM21* mRNA levels are measured by RT-qPCR and cell lysates were subjected to immunoblotting with the indicated antibodies. (F) PC3 cells were treated with PKI-587 (1 µM) for indicated time periods. *TRIM21* mRNA levels are measured by RT-qPCR and cell lysates were subjected to immunoblotting with the indicated antibodies. (G) Human T-ALL and LNCAP prostate cancer cells were treated with PKI-587 (1 µM) for the indicated time periods. *TRIM21* mRNA levels are measured by RT-qPCR and cell lysates were subjected to immunoblotting with the indicated antibodies. (H) Correlations between PI3K pathway activity score and *TRIM21* mRNA expression levels in human thyroid and testicular cancers. Data are presented as fold changes, means ± SD, and compared to WT cells. Each experiment was performed at least three independent times. * p < 0.05, ** p < 0.001, and *** p<0.001, based on Student’s *t*-test. See also Figure S4.

PI3K/AKT signaling also plays similar role in regulating TRIM21 expression and G6PD level in human cancer cells. *TRIM21* mRNA and protein levels were upregulated when WT PTEN, but not phosphatase-dead CS PTEN, was re-expressed in PC3 cells (Figure 4E). PKI-587 treatment led to increased *TRIM21* mRNA and protein levels, accompanied by reduced G6PD and known TRIM21 substrate IRF3 ^29^ protein levels in a time-dependent manner (Figure 4F).

We further extended our studies by analyzing additional prostate cancer and T-ALL lines, all of which carried *PTEN* mutations or deletions ^30, 31^(Supplemental Table S2). Since the tumor suppressor p53 and oncogene c-MYC play critical roles in regulating glucose metabolism^5, 26, 27^, we also included cancer lines with p53 mutations or MYC overexpression to test whether the PI3K-regulated *TRIM21* expression depend on the p53 or MYC status. All human cancer cells tested responded to the PKI-587 treatment in a similar manner, i.e., with a time-dependent increases in *TRIM21* mRNA and protein levels, accompanied by decreased total G6PD levels (Figure 4G), which are independent of p53 and MYC status. Importantly, a strong negative correlation was also found between PI3K pathway activity and *TRIM21* expression in several human cancers (Figure 4H; Spearman’s rank correlation rho was -0.63 and -0.61 respectively, with p value lower than 0.01; Figure S4B). Taken together, these data suggested that the PI3K/AKT pathway negatively regulate the expression of the E3 ligase *TRIM21*, which in turn controls the level and activity of the PPP rate-limiting enzyme G6PD via ubiquitin-mediated degradation *in vivo* and in human cancers.

### PPP activates AKT and support cell growth in the presence of a constitutively activated PI3K pathway

PPP plays an essential role in cell growth even in the presence of a constitutively activated PI3K pathway in PTEN null cells. Blocking PPP by 6-AN significantly inhibited mES colony formation in the presence of a normal concentration of glucose, similar to the effects of glucose starvation and PKI-587 treatment (Figure 5A, left panel). FACS analysis further demonstrated that aforementioned treatments led to similar levels of cell cycle blockage as well as apoptosis (Figure 5A, middle and right panels). Since PI3K/AKT plays central role in cell proliferation and survival, we hypothesized that PPP may feedback regulate PI3K/AKT activity.

**Figure 5.**
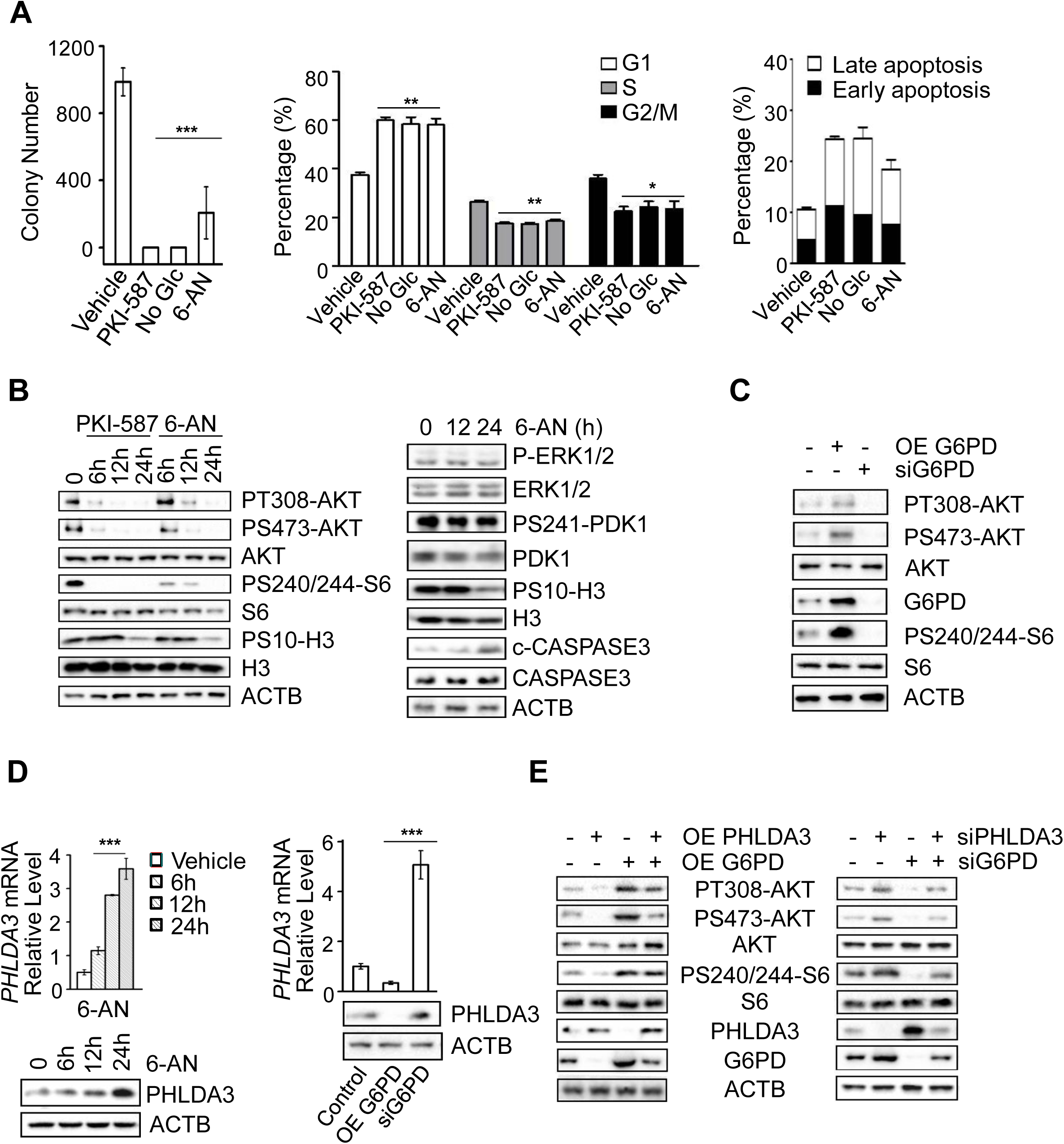
PPP promotes cell growth and AKT activation in the presence of a constitutively activated PI3K pathway by negatively regulating PHLDA3. (A) 6-AN inhibits cell growth, similar to glucose starvation and PKI-587 treatment. *Pten* null mES cells were seeded in 3.5 cm dishes at a density of 2,000 cells/well. Forty-eight hours later, cells were treated with PKI-587 (1 µM), 6-AN (100 nM) or glucose depletion. Colony numbers were counted 7 days later (left panel) while percentages of cells in each phase of the cell cycle and apoptosis were measured by FACS analysis after 24hr of PKI-587 (1 µM) or 6-AN (100 nM) treatment or 12hr in glucose-depleted medium (right panel). (B) 6-AN inhibits AKT activity, similar to the effect of PKI-587 treatment (left panel), but has no effect on ERK and PDK1 activities (right panel). *Pten* null mES cells were treated with PKI-587 (1 µM) or 6-AN (100 nM) for indicated time periods. Cell lysates were subjected to immunoblotting with the indicated antibodies. (C) Overexpression or knock-down of G6PD alters AKT activities in PC3 cells. (D) The levels of the *PHLDA3* mRNA and protein are regulated by 6-AN and *G6PD* activity in PC3 cells. (E) The PPP regulates AKT activity through a PHLDA3-dependent mechanism. *PHLDA3* overexpression abolishes *G6PD* overexpression-induced AKT activation (left) while *PHLDA3* knockdown blocks *G6PD* knockdown-induced AKT inactivation (right). Data are presented as the means ± SD and compared to WT or untreated cells. Each experiment was performed at least three independent times. * p < 0.05, **p < 0.001, and *** p<0.001, based on Student’s *t*-test. See also Figure S5.

Indeed, 6-AN treatment could effectively inhibit AKT phosphorylation and activation of its downstream effector S6, similar to PKI-587 treatment (Figure 5B, left panel). Importantly, associated with 6-AN or PKI-587-mediated AKT inhibition, phospho-H3 levels were significantly decreased while c-CASPASE3 levels were significantly increased (Figure 5B). However, 6-AN treatment had no effect on ERK and PDK1 activities, suggesting an AKT-specific effect (Figure 5B, right panel).

We also tested the association between PPP and AKT activity in *PTEN* null human prostate cancer and T-ALL cell lines. 6-AN treatment substantially decreased P-AKT levels, similar to the effects of the PKI-587 treatment or glucose starvation, accompanied by significant change of apoptosis and cell cycle (Figure S5A and S5B). Therefore, PPP also activates AKT and support cell growth and survival in human cancer cells.

To avoid nonspecific effects of 6AN, we manipulated PPP activity by overexpressing or knocking down its rate limiting enzyme G6PD in the *PTEN* null prostate cancer PC3 cells. G6PD overexpression increased AKT phosphorylation, while *G6PD* knockdown significantly decreased AKT phosphorylation (Figure 5C). Thus, increased PPP metabolism can activate AKT through a positive feedback mechanism even in *PTEN* mutant cells and in the presence of the constitutively activated PI3K pathway.

### PPP control AKT activation by negatively regulating the tumor suppressor PHLDA3

To explore the mechanism of PPP-regulated AKT activation, we conducted RNAseq analysis on 6-AN-treated cells and found that, among those molecules known to negatively regulate AKT activation, the levels of the pleckstrin homology-like domain family A member 3 (*PHLDA3)* mRNA and protein were significantly increased following the 6-AN treatment (Figure 5D, left panel). Similarly, the levels of the *PHLDA3* mRNA and protein were increased upon *G6PD* knockdown and decreased upon *G6PD* overexpression in human *PTEN* null PC3 prostate cancer cells (Figure 5D, right panel), indicating that *PHLDA3* expression is negatively regulated by the PPP.

As a new identified tumor suppressor, PHLDA3 suppresses AKT activation by directly competing with the AKT PH domain for binding to the membrane-associated PIP3, the first step in AKT activation^32^, which may explain the AKT-specific effect observed in our study (Figure 5B). We therefore further determined the functional relevance of PHLDA3 in PPP-regulated AKT activation in *G6PD* overexpressed or knockdown cells. AKT activation was significantly abolished by *PHLDA3* overexpression even in G6PD overexpressed cells (Figure 5E, left panel). *PHLDA3* knockdown, on the other hand, led to increased AKT activation and resistant to G6PD knockdown- or 6-AN-mediated AKT inhibition (Figure 5E, right panel; Figure S5C). Together, these results indicate that PHLDA3 acts downstream of G6PD-regulated PPP pathway and PPP feedback regulate AKT activation by inhibiting the expression of *PHLDA3*.

### PPP metabolites promote AKT activation by negatively regulating PHLDA3 expression

To further investigate how PPP regulates AKT activation, we tested several key PPP metabolites that could be up taken by cells from the media^33, 34^. Addition of R5P or uridine could effectively rescue glucose starvation-induced P-AKT reduction in a concentration-dependent manner, accompanied by decreased PHLDA3 levels (Figure 6A). We then conducted a time course study and found that exogenous R5P and uridine could incorporate into PPP pathway within 10 min (Figure S6A), and significantly down regulate PHLDA3 mRNA and protein levels and activate AKT within 30-60min with or without glucose starvation (Figure 6B and 6C; Figure S6B), suggesting that the PPP-PHLDA3-AKT feedback loop functions under both normal and glucose starved conditions. The effects of R5P and uridine on AKT activation are unlikely due to increased cell proliferation, as H3-S10 phosphorylation levels did not change significantly within the time window tested (Figure 6B). P-ERK and P-PDK1 showed no obvious changes after R5P or uridine addition, further confirming that PPP regulated cell signaling is AKT-specific (Figure S6C and data not shown).

**Figure 6.**
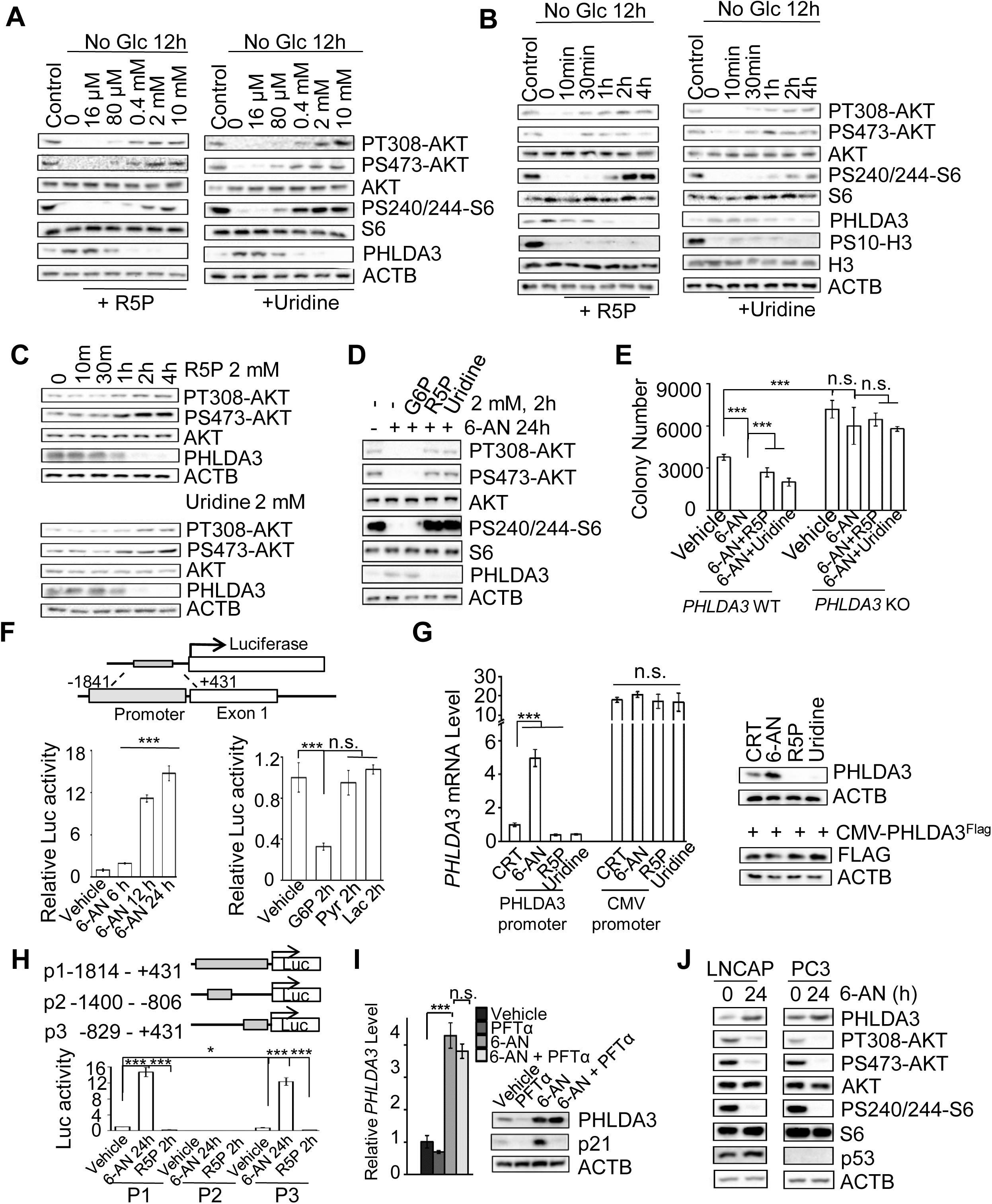
PPP metabolites promote AKT activation by negatively regulating PHLDA3. (A) The addition of the PPP metabolites R5P or uridine blocks glucose starvation-induced PHLDA3 upregulation and AKT inhibition in a concentration-dependent manner. *Pten* null mES cells were treated with various concentrations of R5P (left panel) or uridine (right panel) for 4 h after 12 hours of glucose starvation. Cell lysates were subjected to immunoblotting with the indicated antibodies. (B) The addition of the PPP metabolites R5P or uridine blocks glucose starvation-induced PHLDA3 upregulation and AKT inhibition in a time-dependent manner. *Pten* null mES cells were treated with 2 mM R5P (left panel) or uridine (right panel) for the indicated time periods after 12 hours of glucose starvation. Cell lysates were subjected to immunoblotting with the indicated antibodies. (C) The addition of the PPP metabolites R5P or uridine decreases PHLDA3 level and increases AKT activation in a time-dependent manner under normal growth condition. *Pten* null mES cells were treated with 2 mM R5P (upper panel) or uridine (lower panel) for the indicated time periods. Cell lysates were subjected to immunoblotting with the indicated antibodies. (D) R5P or uridine reverts 6-AN treatment-induced PHLDA3 upregulation and AKT inhibition. *Pten* null mES cells were treated with 6-AN (100 nM) for 24 hours, followed by treatment with 2 mM R5P, uridine and G6P for 2 hours. Cell lysates were subjected to immunoblotting with the indicated antibodies. (E) PHLDA3 knockout blocks the effects of 6-AN, R5P or uridine on colony formation. *PHLDA3* WT and knockout HeLa cells were seeded in 3.5 cm dishes at a density of 2,000 cells/well. Forty-eight hours later, cells were treated with 6-AN (100 nM) in the presence or absence of 2 mM R5P or Uridine. Three days later, colony numbers were counted (E). (F) The PPP pathway controls PHLDA3 expression by regulating its promoter activity. Upper panel, a schematic map of PHLDA3 promoter-luciferase reporter construct. PC3 cells were transfected with the PHLDA3-Luciferase reporter plasmid. Twenty-four hours later, cells were treated with 6-AN (100 nM) for the indicated time periods (lower left panel) or incubated with 2 mM G6P, pyruvate, or lactate for 2hr, respectively (lower right panel). The cell lysates were harvested for luciferase activity assay. (G) PHLDA3 driven by an exogenous promoter fails not response to 6-AN treatment nor R5P and uridine supplements. PC3 cells were transfected with or without CMV driven FLAG-PHLDA3 expression plasmid. Then cells were treated with 6-AN (100 nM) for 24 hours or 2 mM R5P or uridine for 2 hours. Cell lysates were subjected to RNA extraction for RT-qPCR or immunoblotting with the indicated antibodies. (H) Truncation analysis further narrowed down the *PHLDA3* promoter region necessary for PPP regulation to p3 (-829 - +431). (I-J) PPP regulates PHLDA3 expression and AKT activation independent of the p53 status. *Pten* null mES cells were treated without or with PFTα (5 μM) or 6-AN (100 nM) for 24 h without and with PFTα (5 μM). *Phlda3* mRNA levels (left panel) and protein levels were measured (I). PTEN null LNCAP (*p53* WT) and PC (*p53* null) cells were treated with 6-AN (100 nM) for 24 h. Cell lysates were subjected to immunoblotting with the indicated antibodies. Data are presented as the means ± SD and compared to WT or untreated cells. Each experiment was performed at least three independent times. * p < 0.05, ** p < 0.001, and *** p<0.001, based on Student’s *t*-test. See also Figure S5.

Since parts of glycolysis and PPP are interconvertible, we further tested the requirement of PPP for AKT activation by adding those glycolytic metabolites that can or cannot convert to PPP. PPP upstream metabolite G6P could activate AKT, while glycolytic downstream products pyruvate and lactate, which cannot convert to PPP metabolism, did not exert a significant effect on P-AKT and PHLDA3 levels within the time window tested (Figure S6D). In addition, treating cells with PPP inhibitor 6-AN or glycolysis inhibitor 2-DG had no effect on R5P or uridine-induced PHLAD3 downregulation and AKT activation but 6-AN could block G6P-induced AKT activation (Figure 6D; Figure S6E), suggesting a PPP-specific regulatory mechanism. Notably, the concentration of R5P and uridine were decreased below physiological levels after 6-AN treatment or glucose starvation, while R5P or uridine supplement was able to rescue their levels within 10 minutes (Figure S6F), indicating that the concentrations of PPP metabolites are important for PHLDA3-regulated AKT inhibition.

6-AN induced growth inhibition was PHLDA3-dependent and colonogenic growth inhibition caused by 6-AN treatment could be rescued by R5P or uridine only in *PHLDA3* WT cells (Figure 6E). Therefore, PHLDA3 acts down stream of PPP metabolism pathway and negatively regulates AKT activity and cell proliferation.

### PPP metabolites negatively regulate *PHLDA3* expression independent of p53

To further investigate the mechanism underlying PPP-regulate *PHLDA3* transcription, we made a *PHLDA3-luciferase* reporter construct containing -1841 - +431 region of the human *PHLDA3* gene (Figure 6F, upper panel). When transfected into PC3 cells, the reporter construct showed similar kinetics of response to 6-AN treatment and R5P or uridine supplement as compared to the endogenous gene (Figure 6F, lower left panel; Figure S6G). G6P showed similar inhibition, while pyruvate and lactate had no obvious effect on PHLDA3 promoter activity (Figure 6F, lower right panel). Importantly, *PHLDA3* driven by an exogenous promoter did not response to 6-AN treatment nor R5P and uridine supplements (Figure 6G).

Truncation analysis further narrowed down the minimal region between -829 and +431 to be necessary for PPP inhibition (Figure 6H). *PHLDA3* expression is known to be regulated by the tumor suppressor p53^32, 35^. The region that p53 binds to the *PHLDA3* promoter overlaps with the minimal region required for PPP-regulated *PHLDA3* expression, which prompted us to investigate whether PPP-regulated *PHLDA3* expression is p53-dependent. For this, we tested the effect of p53 inhibitor PFTα on 6-AN-induced *PHLDA3* expression. Although we observed a slightly reduced *Phlda3* level when we treated the *Pten* null mES cells with the p53 inhibitor PFTα, we did not observe any significant difference in 6-AN treatment-induced *Phlda3* expression without or with PFTα (Figure 6I). Importantly, *PTEN* null *p53* WT LNCAP and *PTEN* and *p53* null PC3 human prostate cancer lines showed similar responses to 6-AN treatment (Figure 6J). These results suggest that PPP-regulated *PHLDA3* expression is p53-independent although further study is needed to identify key transcription factor involved in PPP -regulated *PHLDA3* promoter activity.

### PPP controls cell proliferation and survival *in vivo* in a PHLDA3-dependent manner

As a very small percentage of glycolytic intermediates are shunted through PPP in normal cells, the dependency of PTEN null cells on PPP may represent an attractive therapeutic opportunity. To test this hypothesis, we treated the *Pten* null prostate cancer and T-ALL models with 6-AN. 6-AN treatment significantly increased the levels of *Phlda3* mRNA and protein, accompanied by substantial decreases in the levels P-AKT and P-S6 as well as the PPP metabolic intermediates (except for 6PG since 6-AN blocks the first two steps of the PPP) (Figure 7A, B, C). As a consequence of the decreased AKT activity, cell proliferation was inhibited, as evidenced by decreased phospho-H3 levels, while cell death was increased, as evidenced by increased cleaved CASPASE3 levels (Figure 7A), indicating that PTEN null cancer cells are sensitive to anti-PPP therapies even in the presence of a hyperactivated PI3K pathway.

**Figure 7.**
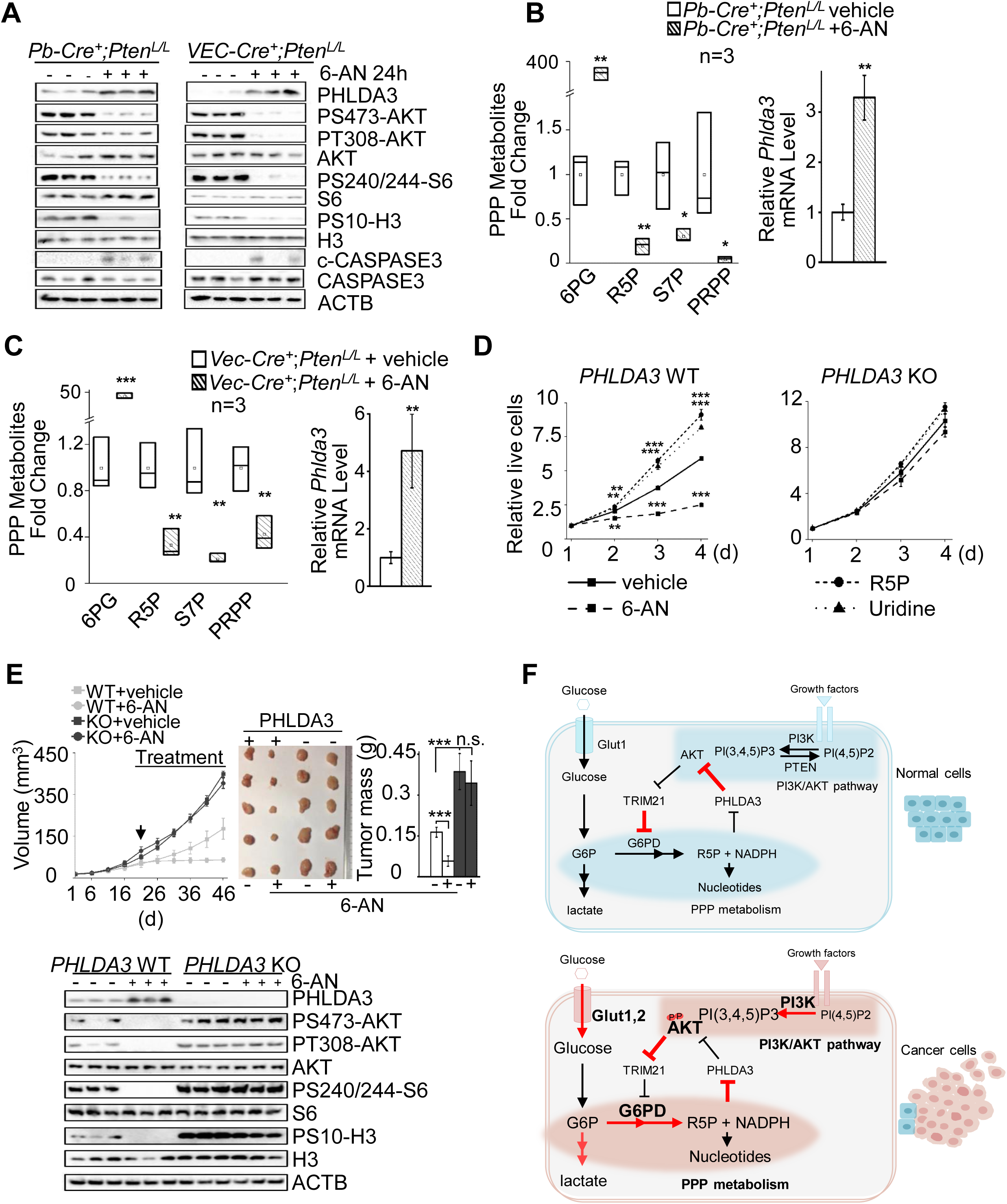
PPP controls cell proliferation and survival *in vivo* in a PHLDA3-dependent manner. (A-C) The effects of PPP inhibition in the *Pten* null prostate cancer and T-ALL mouse models. Twenty-four hours 6-AN (15mg/kg) treatment leads to increased *Phlda3* mRNA and protein levels and cleaved caspase 3 levels, decreased P-AKT, P-H3 and PPP metabolites. Metabolites in PPP were extracted from the anterior lobes of *Pb-Cre^−^;Pten^L/L^*and *Pb-Cre^+^;Pten^L/L^* prostates or thymus of *VEC-Cre^−^;Pten^L/L^* and *VEC-Cre^+^;Pten^L/L^* mice without or with 6-AN treatment, then measured by LC-MS and normalized to protein concentration (left panel), mRNA were detected by RT-PCR and cell lysates were subjected to immunoblotting with the indicated antibodies. (D) PHLDA3 knockout blocks the effects of 6-AN, R5P or uridine on cell growth *in vitro*. *PHLDA3* WT and knockout cells were seeded in 96-well plates at a density of 1,000cells/well, respectively. Forty-eight hours later, cells were treated with 6-AN (100 nM) or 2 mM R5P or uridine. Four days later, the cell viability was detected by CCK-8 assay (E) *PHLDA3* knockout blocks 6-AN-induced cell growth *in vivo*. Equal numbers of *PHLDA3* WT and *PHLDA3* knockout cells were implanted onto the bilateral flanks of nude mice. Tumor volumes and weights were measured and presented (upper panel). Cell lysates from the xenograft tumors were subjected to immunoblotting with the indicated antibodies (lower panel). (F) Schematic representation of the cross-talk between the PI3K/AKT pathway and PPP metabolism. TRIM21 and PHLDA3 are two negative regulators identified in this study. TRIM21 and PHLDA3 play essential roles in regulating the AKT-TRIM21-G6PD (PPP) and PPP-PHLDA3-AKT feedback loops, respectively, in this signaling pathway-metabolism pathway crosstalk. Data are presented as the means ± SD and compared to WT or vehicle-treated cells. Each experiment was performed at least three independent times. * p < 0.05, ** p < 0.001, and *** p<0.001, based on Student’s *t*-test. (H) TRIM21 and PHLDA3 Negatively Regulate the Cross-Talk between the PI3K/AKT Pathway and PPP Metabolism See also Figure S7.

The response of cells to PPP inhibition is PHLDA3 dependent. *PHLDA3* knockout cells have hyperactivated AKT and did not response to the inhibition effects of 6-AN on AKT phosphorylation and cell growth (Figure 7D; Figure S7A). As PHLDA3 acts downstream of PPP, *PHLDA3* knockout cells did not response to the growth promotion effect of R5P or uridine supplement either, when compared to *PHLDA3* WT cells (Figure 7D). We also tested the effects of 6-AN treatment on *PHLDA3* WT- and knockout-derived xenograft models *in vivo*. As shown in Figure 7E, 6-AN treatment could not inhibit *PHLDA3* knockout xenograft tumor growth *in vivo* (Figure 7E, upper panels) due to the constitutively activated AKT pathway (Figure 7E, lower panel). These results suggest that cancers with *PHLDA3* loss may be resistant to anti-PPP inhibitor treatment.

## DISCUSSION

Most cancer metabolism studies are focused on how oncogene and tumor suppressor genes, as well as their controlled signaling pathways, regulate cell metabolism. Our study suggests that metabolic pathway is not a passive recipient of oncogenic signals, rather it feedback regulates oncogenic signals. In normal cells, PTEN restrains PI3K/AKT activity and promotes TRIM21-mediated G6PD degradation, and consequently lower down PPP activity and cell proliferation. PPP metabolites, on the other hand, keep AKT activity in check by negatively regulating the expression of its inhibitor *PHLDA3*. Loss of PTEN tumor suppressor or activation of PI3K/AKT pathway in cancer cells enhances G6PD activity and PPP metabolism by inhibiting TRIM21. Increased PPP metabolites, in turn, blocks *PHLDA3* expression and reinforce AKT activation. The reciprocal cross-talks between oncogenic signaling and metabolic pathways, therefore, may serve as a positive enforcement for cancer metabolic reprogram and ensure uncontrollable cancer growth. The negative regulators in these cross-talks, such as TRIM21 and PHLDA3 discovered in this study, may contribute to cancer development and treatment resistance (Figure 7F).

By studying isogenic mES cells, we found that PTEN controls glucose metabolism mainly through its lipid phosphatase activity by antagonizing the PI3K/AKT pathway. PTEN loss or PI3K activation not only promotes the “Warburg effect” by increasing the level of a glucose transporter and the rate of glycolysis, but also decouples glycolysis and the TCA cycle and diverts glycolytic intermediates to the branching metabolism pathways PPP (Figure 1).

The effect of PI3K/AKT activation on the PPP metabolic pathway observed in this study differs from that of oncogene KRAS^36^. While both the oxidative and nonoxidative arms of PPP, together with PPP-associated nucleotides and NADPH production, are upregulated in the *Pten* null cells (Figure 1), only the nonoxidative arm is upregulated in the *Kras* pancreatic cancer model^36^. These differences are most likely mediated by the mechanisms by which PI3K/AKT and KRAS control the PPP. As described in our study, PI3K/AKT controls the activity of G6PD, which regulates the oxidative arms of PPP and determines the speed of glycolytic intermediate flux into PPP (Figure 2). AKT is also known to modulate TKT activity, a key enzyme in the nonoxidative PPP^37, 38^. KRAS, on the other hand, controls the expression of enzymes involved in the nonoxidative arm of the PPP^36^. As the oxidative arm of the PPP is irreversible and important for nucleotide and NADPH production^38^, the difference between the effects of PI3K/AKT and KRAS signaling on the PPP metabolic pathway may explain the increased oxidative stress experienced by KRAS tumors.

Ubiquitylation and proteasome-mediated degradation are instrumental in regulating protein functions. In our study, PTEN loss of function or PI3K/AKT activation prolonged the G6PD half-life by preventing its ubiquitin-mediated degradation – a novel mechanism of G6PD regulation (Figure 2). We also identified TRIM21 as the E3 ligase for G6PD, which is transcriptionally downregulated by PI3K/AKT activation (Figures 3 and 4). Our analysis demonstrates TRIM21 expression levels are negatively correlated with PI3K activity in various human cancers, implying that TRIM21 may act as a potential tumor suppressor. TRIM21 can also promote phosphofructokinase 1 (PFK1) degradation^28^. Therefore, TRIM21 may act at multiple nodes to control normal cell metabolism and cancer metabolic reprogramming.

In addition to the mechanisms by which PI3K/AKT regulated glycolysis and PPP, we also discovered a novel pathway by which PPP metabolites feedback positively regulates AKT activation, even in the presence of a constitutively activated PI3K pathway (Figures 5-7). It has been reported that intracellular nutrient or energy information can transmit to mTORC1^39–41^ and AMPK^42, 43^. Here we identified a new mechanism by which PPP metabolites promote AKT activation, cell proliferation and survival via inhibition of a tumor suppressor PHLDA3 (Figures 5 and 6). *PHLDA3*, encoding a PH domain-only protein, is a negative regulator of AKT activation by competing binding to PIP3^32^. *PHLDA3* loss of heterozygosity has been reported in human lung and neuroendocrine pancreatic cancers^32, 35^. We found that, in PTEN null mES and *in vivo* cancer models, the abundant PPP metabolites can restrain *PHLDA3* expression, which in turn activates AKT and further promotes cell growth. Importantly, a similar control mechanism is also present in human prostate and T-ALL lines.

Different from previously reported p53-regulated *PHLDA3* expression^32^, our study indicates that PPP-regulated *PHLDA3* expression is p53-independent. PPP-regulated cell signaling is also AKT-specific as it has no effect on the activities of either the AKT upstream PDK-1 or parallel pathway ERK (Figure 6). Therefore, our study discovered a novel reciprocal feedback control mechanism between the PI3K/AKT pathway and the PPP metabolism (Figure 7F), which not only orchestrates the partitioning of glycolytic intermediates to branching metabolic pathways to meet the needs for rapid tumor cell growth but also reinforces PI3K/AKT activation and aerobic glycolysis for cancer metabolic reprogramming.

As a very small percentage of glucose is shunted through the PPP in normal cells, the dependency of PTEN null cells on the PPP and the reciprocal cross-talk between the PI3K/AKT pathway and the PPP may represent an attractive therapeutic opportunity. As a proof of principle study, we revealed that PPP inhibition impeded AKT activation and the proliferation and survival of PTEN null mES cells and human cancer cells, as well as *in vivo* cancer models, indicating that PTEN null cancer cells may be more vulnerable to anti-PPP therapy, even in the presence of a hyperactivated PI3K pathway. We also demonstrated that the effects of PPP inhibition on AKT activation and cell proliferation are PHLDA3-dependent and predict that the PHLDA3 status in human cancers may dictate their response to anti-PPP treatments.

## AUTHORS’ CONTRIBUTIONS

J.C., Y.H. and H.W. conceived the project. J.C. and Y.H. performed the majority of the experiments; S.W. initiated PKM2 studies; J.J. generated *Pten* mutant mES lines; M.W., L.T., and F.C. conducted bioinformatics analysis; L.Z., M.W., W.Y. and Y.W. contributed to experiments involving *in vivo* mouse models; all under the supervision of H.W. XH.Z and Y.Y. conducted glycolytic/TCA and PPP metabolic measurements under the supervision of XX.Z; P.J. provided essential materials and suggestions. J.C., Y.H. and H.W. wrote the manuscript with help from all authors.

## ACKNOWLEDGMENTS

We appreciate Dr. Utpal Banerjee for sharing unpublished data; Li Chen in Dr. Joshua D. Rabinowitz laboratory and Dr. Qunying Lei, and members of our laboratory for helpful comments and insightful suggestions. We also thank the Metabolomics Facility and the Functional Analysis Core Facility of the National Center for Protein Sciences Beijing (Peking and Tsinghua Universities) for technical assistance. This study was supported by the Peking-Tsinghua Center for Life Sciences and Beijing Advanced Innovation Center for Genomics at Peking University to HW.

---

**Figure S1.**
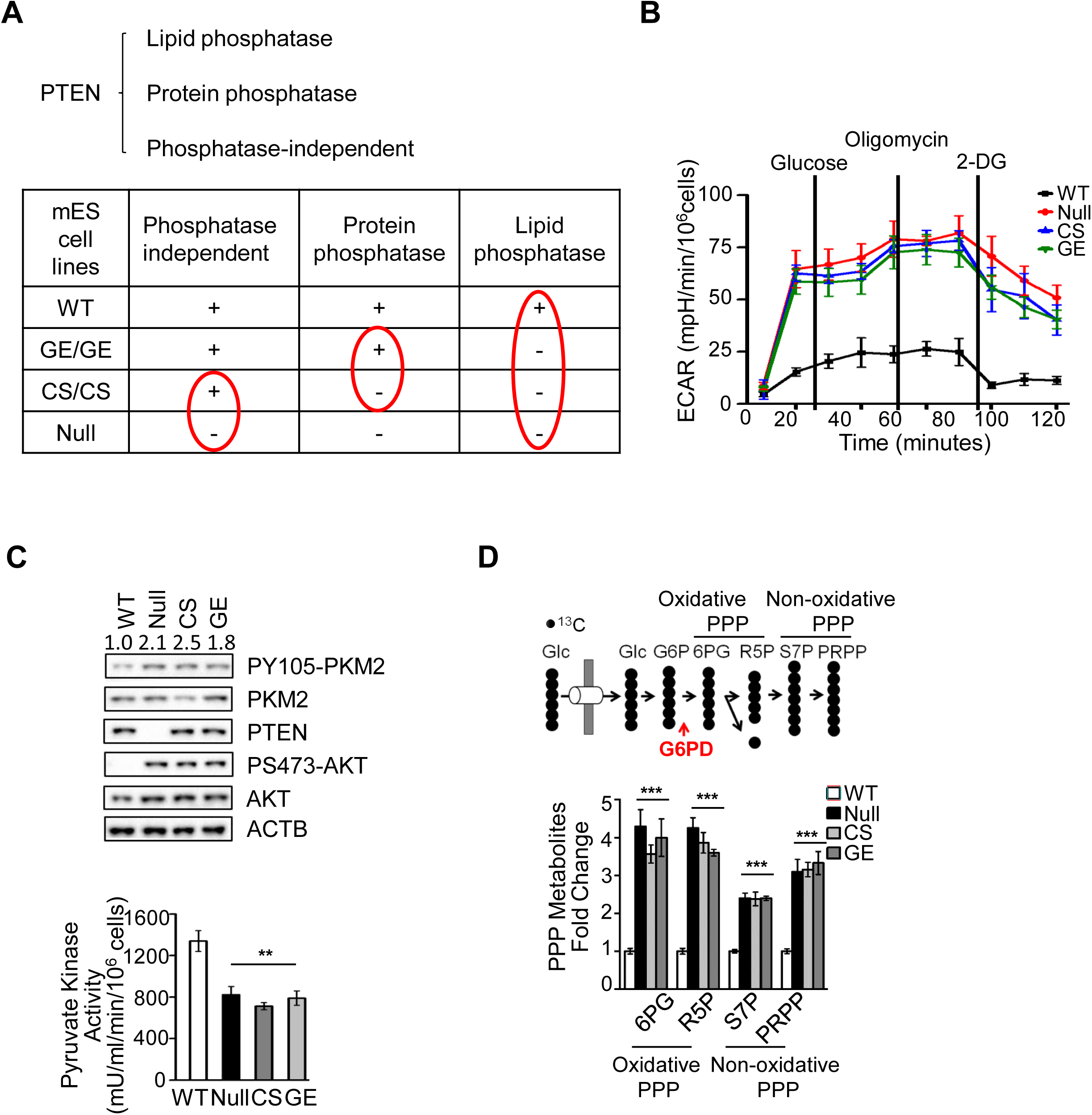
PTEN loss of function diverts glycolytic intermediates to branching pathways by activating PI3K. (A) Upper panel, major biological functions of PTEN; lower panel, genetic separation of the major biological functions of PTEN by site-specific knock-in mutations and deletions. (B) PTEN loss of function or PI3K activation increases ECAR rates. *Pten* WT, null, CS and GE mES cells were cultured without glucose for 1 hour before the sequential addition of glucose (80 mM), oligomycin (2 mM), and 2-DG (50 mM). ECAR was calculated as the rate of PH change per minute per 2×10^4^ cells. (C) Loss of PTEN lipid phosphatase activity increases PY105-PKM2 (upper panel), and decreases PKM2 activity (lower panel). (D) Upper panel, a schematic illustrating [U-^13^C] glucose tracing into the PPP; lower panel, levels of labeled PPP metabolites are increased in *Pten* null, CS, and GE mES cells compared to WT cells. Cell extracts were prepared and analyzed using LC-MS. Data are presented as fold changes, means ± SD, and compared to WT cells. Each experiment was performed at least three independent times. * p < 0.05, ** p < 0.001, and *** p<0.001, based on Student’s *t*-test.

**Figure S2.**
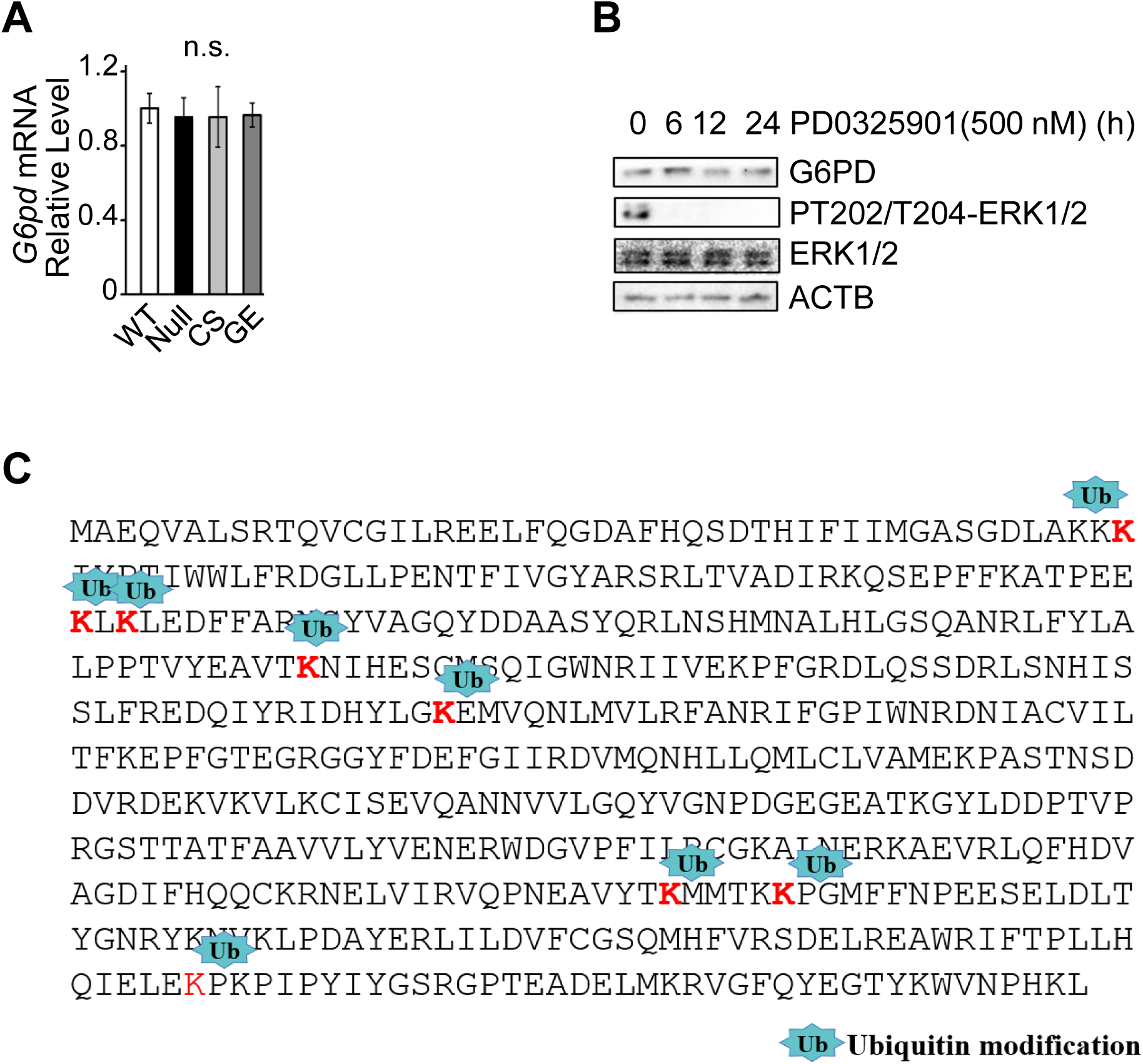
The PI3K/AKT pathway induces the PPP by stabilizing the rate-limiting enzyme G6PD. (A) *G6pd* mRNA levels are not changed in *Pten* WT, null, CS and GE mES cells. (B) Levels of the G6PD protein remain unchanged in PC3 cells treated with PD0325901 (500 nM) for the indicated time periods. (C) The potential G6PD ubiquitination-modified sites identified by affinity MS. Data are presented as fold changes, means ± SD, and compared to WT cells. Each experiment was performed at least three independent times. * p < 0.05, ** p < 0.001, and *** p<0.001, based on Student’s *t*-test.

**Figure S3.**
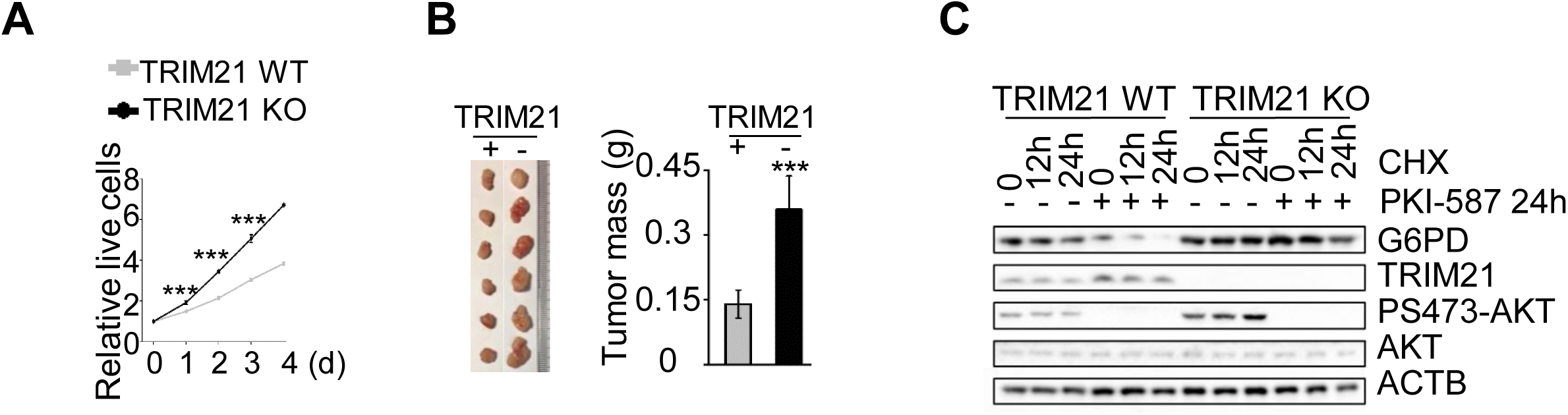
TRIM21 is the E3 ligase responsible for PI3K/AKT- regulated G6PD stability. (A) *TRIM21* knockout leads to increased cells growth *in vitro*. Equal numbers of *TRIM21* WT and knockout cells were seeded in 96 well plates at 2,000 cells/well. Relative cell viabilities were tested using CCK8 kit and presented as growth curves. (B) *TRIM21* knockout leads to increased cells growth *in vivo*. Equal numbers of *TRIM21* WT and knockout cells were implanted onto the bilateral flanks of nude mice. Tumor mass were measured and calculated. (C) TRIM21 controls G6PD stability. Half-lives of G6PD in *TRIM21* WT and knockout cells were measured after vehicle or PKI-587 (1 µM) treatment for 24 h with CHX (100 µg/ml) treatment for the indicated times. Cell lysates were subjected to Western blot analysis using the indicated antibodies. Data are presented as fold changes, means ± SD, and compared to WT cells. Each experiment was performed at least three independent times. * p < 0.05, ** p < 0.001, and *** p<0.001, based on Student’s *t*-test.

**Figure S4.**
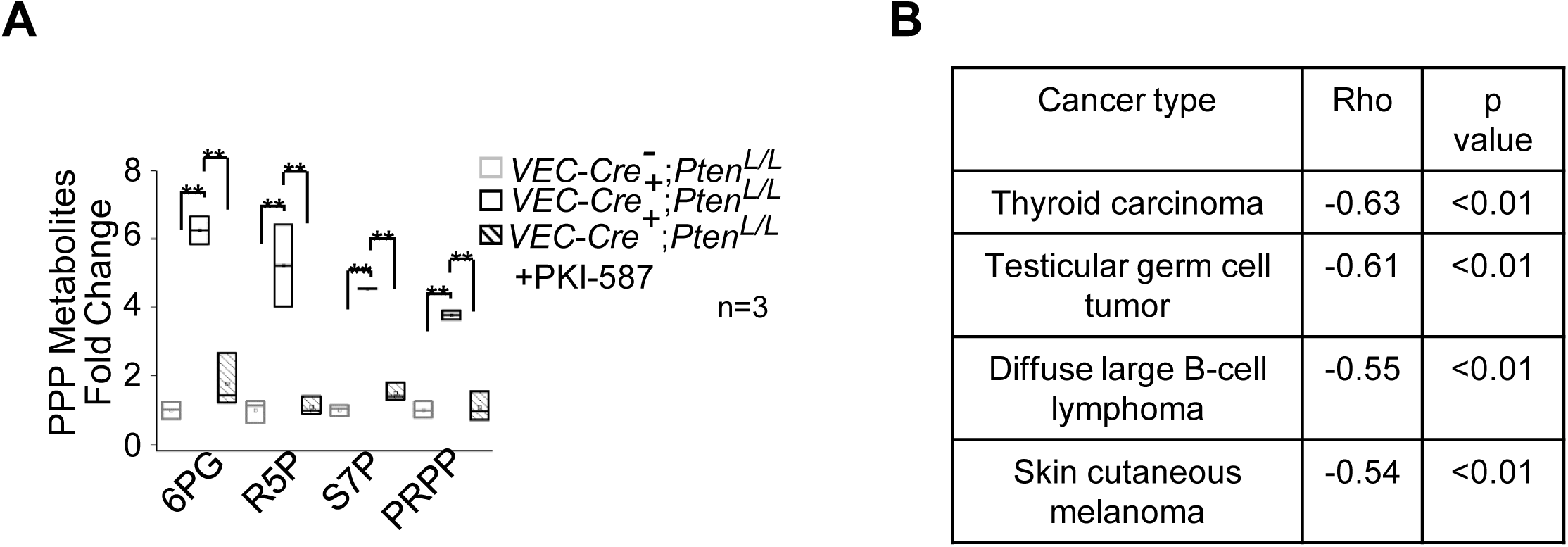
PI3K/AKT regulates TRIM21 and G6PD *in vivo* and in human cancers. (A) PI3K regulates the PPP metabolic pathways in *Pten* null T-ALL mouse model. (B) Correlations between PI3K pathway activity score and TRIM21 mRNA expression levels in human cancers (based on TCGA data). Cell extracts were prepared and analyzed using LC-MS. Data are presented as fold changes, means ± SD, and compared to WT cells. Each experiment was performed at least three independent times. * p < 0.05, ** p < 0.001, and *** p<0.001, based on Student’s *t*-test.

**Figure S5.**
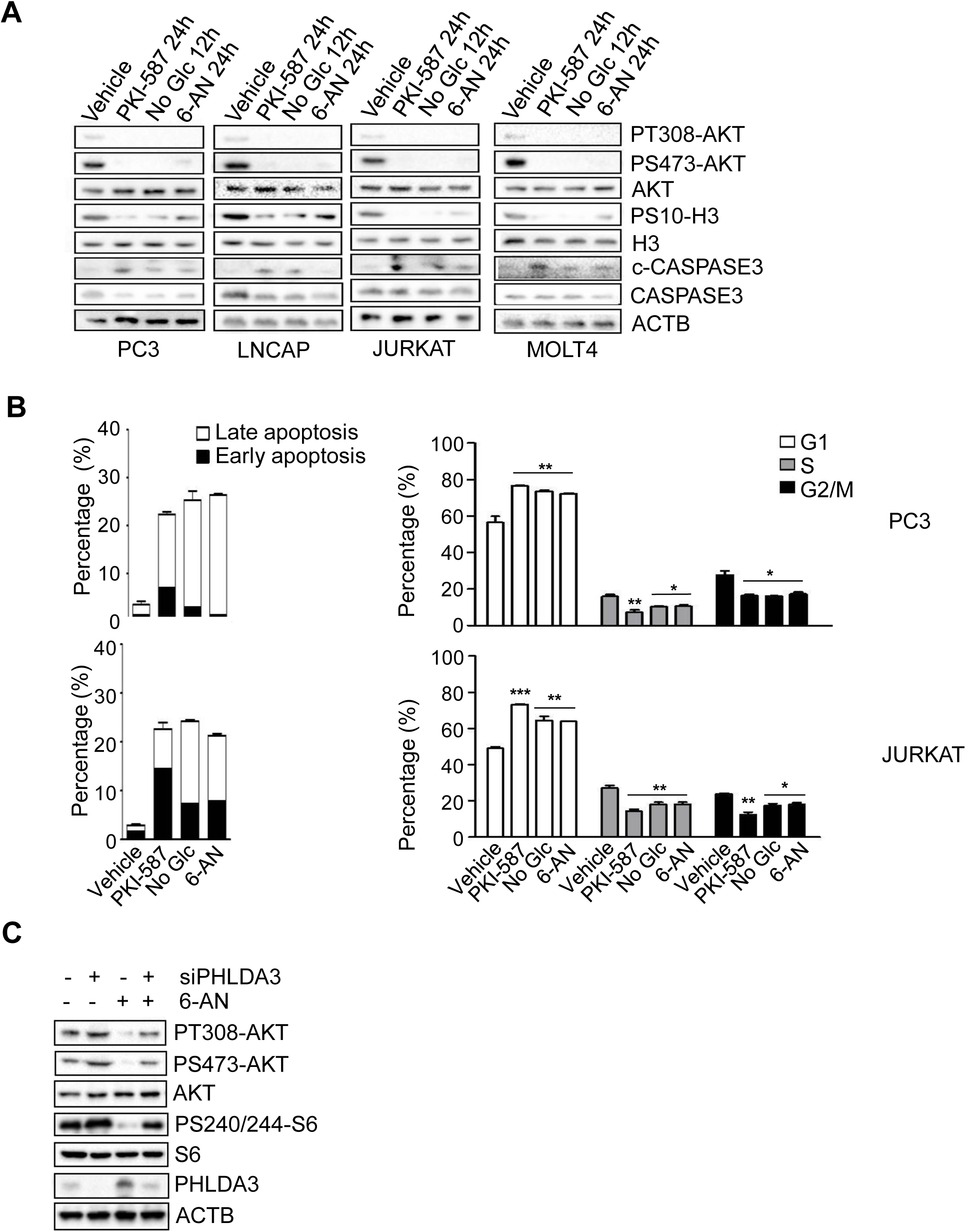
PPP promotes cell growth and AKT activation in the presence of a constitutively activated PI3K pathway by negatively regulating PHLDA3. (A-B) PPP activates AKT and support cell growth in human cancer cell lines. PC3, LNCAP, JURKAT and MOLT4 cells were treated with PKI-587 (1 µM) or 6-AN (100 nM) for 24 h or cultured in glucose-depleted medium for 12 h. Cell lysates were subjected to immunoblotting with the indicated antibodies (A). For PPP regulated cell proliferation and survival, PC3 and JURKAT cells were treated with PKI-587 (1 µM) or 6-AN (100 nM) for 24 h or cultured in glucose-depleted medium for 24 h. The percentages of apoptotic cells and cells in each phase of the cell cycle were determined by FACS analysis. (C) *PHLDA3* knockdown blocks 6-AN (100 nM) treatment-induced AKT inactivation in PC3 cells. Data are presented as fold changes, means ± SD, and compared to WT cells. Each experiment was performed at least three independent times. * p < 0.05, ** p < 0.001, and *** p<0.001, based on Student’s *t*-test. See also Supplemental Table 2.

**Figure S6.**
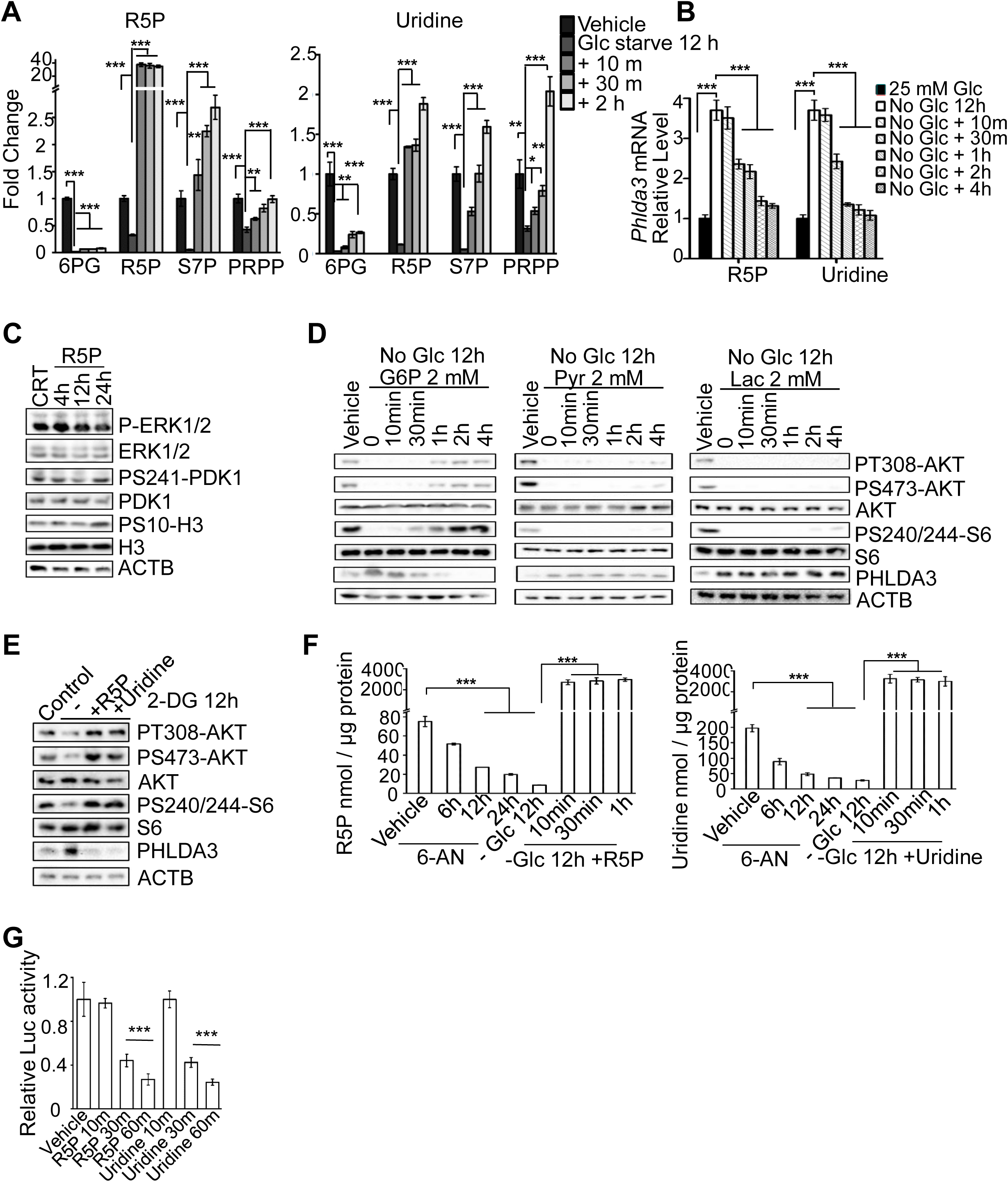
PPP metabolites promote AKT activation by negatively regulating PHLDA3. (A) Exogenous R5P or uridine could immediately incorporate into PPP metabolites. PC3 cells were treated with 2 mM R5P (left panel) or uridine (right panel) for the indicated time periods after 12 hours of glucose starvation. Then the PPP metabolites were extracted from cell lysates and measured. (B) R5P and uridine inhibits *PHLDA3* mRNA expression. PC3 cells were treated with R5P (2 mM) or uridine (2 mM) for indicated time periods after 12 hours of glucose starvation and subjected to RT-PCR. (C) R5P treatment has no effect on ERK and PDK1 pathway. *Pten* null mES cells were treated with R5P (2 mM) for the indicated time periods. Cell lysates were subjected to immunoblotting with the indicated antibodies. (D) The effects of G6P (left penal), pyruvate (middle penal) and lactate (right penal) on AKT activation and PHLDA3 protein levels. *Pten* null mES cells were treated with 2 mM G6P, 10 mM pyruvate, or 25 mM lactate for the indicated times after 12 hours of glucose starvation. Cell lysates were subjected to immunoblotting with the indicated antibodies. (E) 2-DG treatment could not diminish R5P and uridine-induced AKT activation. *Pten* null mES cells were treated with 2 mM R5P or uridine for 2 h after 12 hours of 2-DG (10 mM) treatment. Cell lysates were subjected to immunoblotting with the indicated antibodies. (F) The endogenous R5P and uridine levels were down-regulated by 6-AN treatment or glucose starvation while exogenous R5P or uridine supplement can immediately increase their concentrations above normal endogenous levels. *Pten* null mES cells were treated with 6-AN for indicated time periods or glucose starvation for 12 h then 2 mM R5P or uridine supplement was added to the media for the indicated time periods before measurements. (G) The PHLDA3 promoter-luciferase construct responded to R5P and uridine supplement in a time-dependent manner. PC3 cells were transfected with the reporter construct and 24 h later supplemented with R5P (2 mM) or uridine for 1 h. The cell lysates were harvested for luciferase assay. Cell extracts were prepared and analyzed using LC-MS. Data are presented as fold changes, means ± SD, and compared to WT cells. Each experiment was performed at least three independent times. * p < 0.05, ** p < 0.001, and *** p<0.001, based on Student’s *t*-test.

**Figure S7.**
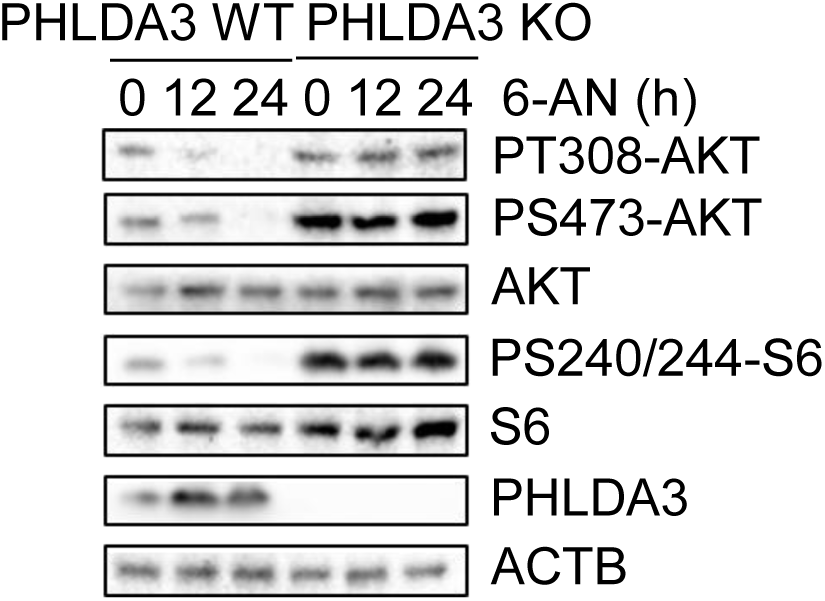
PPP controls cell proliferation and survival *in vivo* in a PHLDA3-dependent manner. The effects of 6-AN treatment-induced PHLDA3 upregulation and AKT inhibition are blocked by PHLDA3 knockout.

**Supplement Table 1:** A list of inhibitors used in this study.

| Drug | Target | IC<sub>50</sub> |
| --- | --- | --- |
| PKI-587 | PI3K/mTOR | PI3Kα=0.4 nM; PI3Kβ, $\delta$, $\gamma$>0.4 nM; mTOR=1.0 nM. |
| BAY1082439 | PI3K | PI3Kα=5 nM; PI3Kβ=15 nM; PI3Kδ=1 Nm; PI3Kγ=52 nM. |
| GDC-0068 | Pan-AKT | Akt1=5 nM; Akt2=18 nM; Akt3=8 nM |
| GDC-0980 | PI3K/mTOR | PI3Kα=5 nM; PI3Kβ=27 nM; PI3Kδ=7 Nm; PI3Kγ=14 nM; mTOR=17 nM. |
| Rapamycin | mTOR | mTOR<0.1 nM |
| PD0325901 | MEK | 0.33 nM |
| 6-Aminonicotinamide<br>(6-AN) | G6PD, PGD | Ki =0.46 μM |

**Supplement Table 2:** A list of human cancer cell lines used in this study.

| Cancer type | Prostate cancer |  | T-ALL |  |  |  |  |  |
| --- | --- | --- | --- | --- | --- | --- | --- | --- |
| Cell line | LNCAP | PC3 | CEM | MOLT3 | MOLT4 | MOLT16 | KE-37 | JURKAT |
| PTEN | Mut | Null | Null | Null | Null | Null | Null | Null |
| p53 | WT | Null | Null | WT | Null | Mut | Null | Null |
| MYC | WT | WT | WT | WT | WT | TCRα-c-myc | TCRα-c-myc | WT |

## EXPERIMENTAL MODEL AND PROCEDURES

### Cell Lines

*Pten* WT, null and mutant (CS and GE) mES cells were adapted to feeder-free culture conditions and maintained in Knock-Out DMEM (Gibco) supplemented with 15% fetal bovine serum (HyClone), 1000 U/ml leukemia inhibitory factor (LIF), 2 mM L-glutamine, 100 U/ml penicillin/streptomycin, 0.1 mM nonessential amino acids (NEAA, Gibco), and 100 µM β-mercaptoethanol. Media were replaced every day and cells were passaged every 3 days. All experiments were performed on mES cells within 15 passages. Glucose-depleted medium for mES cells was purchased from Gibco. T-ALL cells (JURKAT, KE-37, CEM, MOLT3, MOLT4 and MOLT16) were cultured in RPMI-1640 (Gibco) containing 10% fetal bovine serum and 100U/ml penicillin/streptomycin. Human prostate cancer cells (PC3 and LNCAP) and HEK293T cells were maintained in DMEM (Gibco) containing 10% fetal bovine serum and 100U/ml penicillin/streptomycin. PC3 WT/CS-PTEN-inducible cells were maintained in DMEM containing 10% tetracycline-free fetal bovine serum and 100 U/ml penicillin/streptomycin and induced by adding 2 μg/ml doxycycline to the medium for the indicated times to induce PTEN expression.

Glucose-depleted media for T-ALL and prostate cancer cells were purchased from Gibco. *TRIM21*-knockout and WT A549 cells, *PHLDA3*-knockout and WT A HeLa cells were generously provided by Prof. Wensheng Wei and cultured in DMEM containing 10% fetal bovine serum and 100U/ml penicillin/streptomycin.

### Mice

The generation of the *Pb-Cre+;Pten^loxP/loxP^*prostate cancer model and *VEC-Cre^+^*;*Pten^loxP/loxP^* T-ALL model has been described previously^22, 24^. Mouse genotypes were verified by PCR with primers for *Pten* and *Cre*^24^. Prostate and thymic tissues were isolated for metabolite measurements, Western blotting and mRNA analysis.

### Key resources table

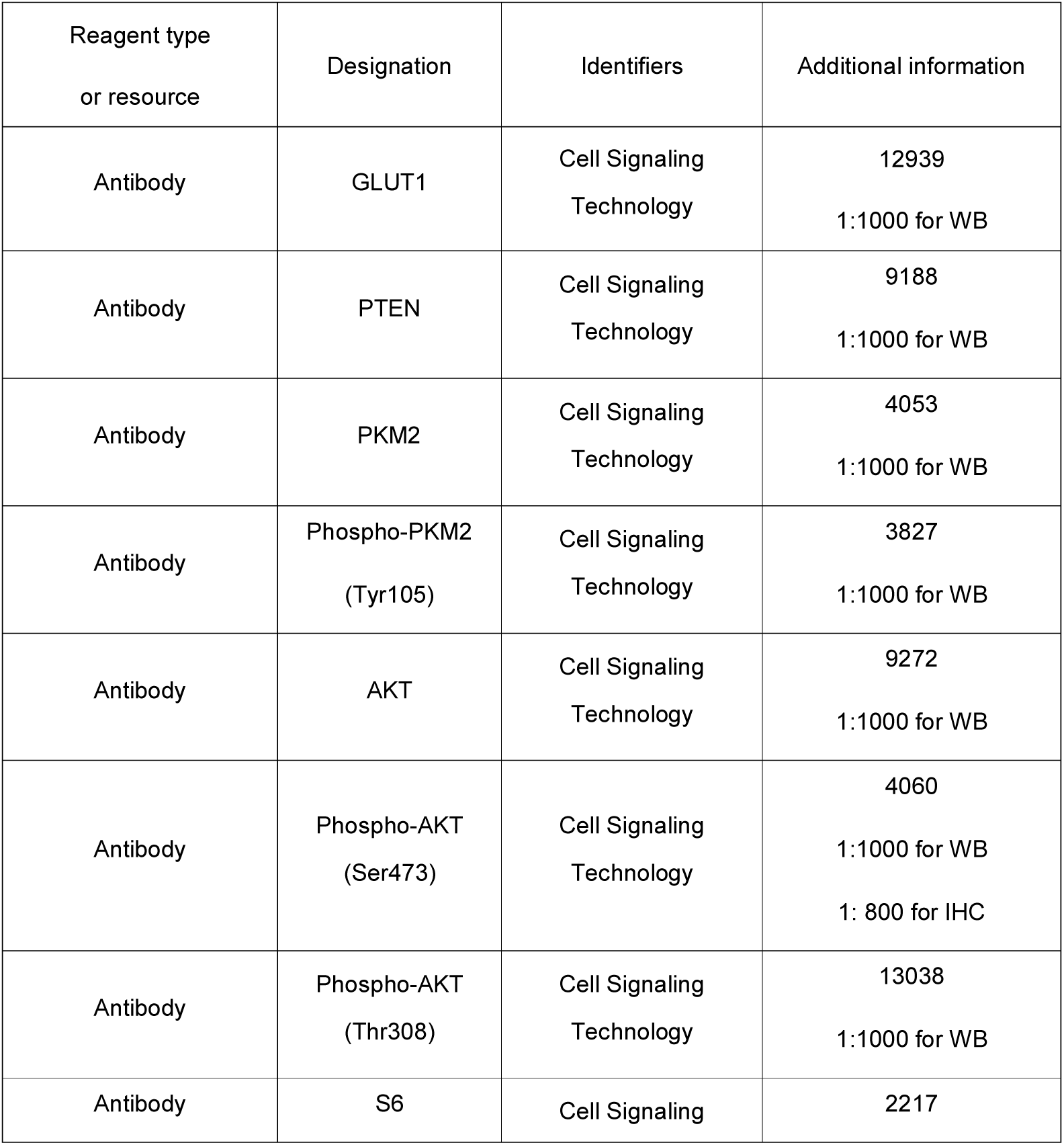
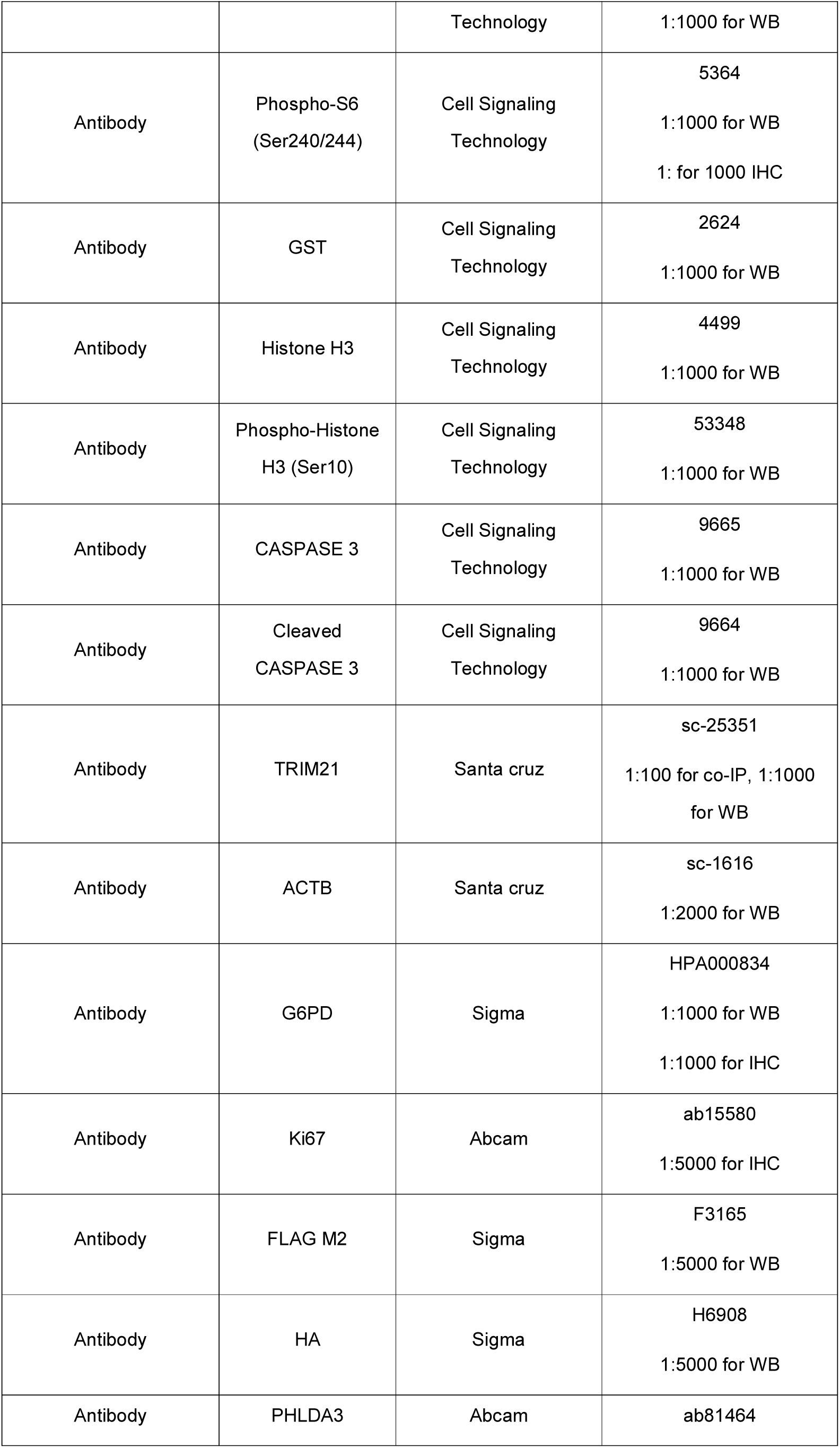
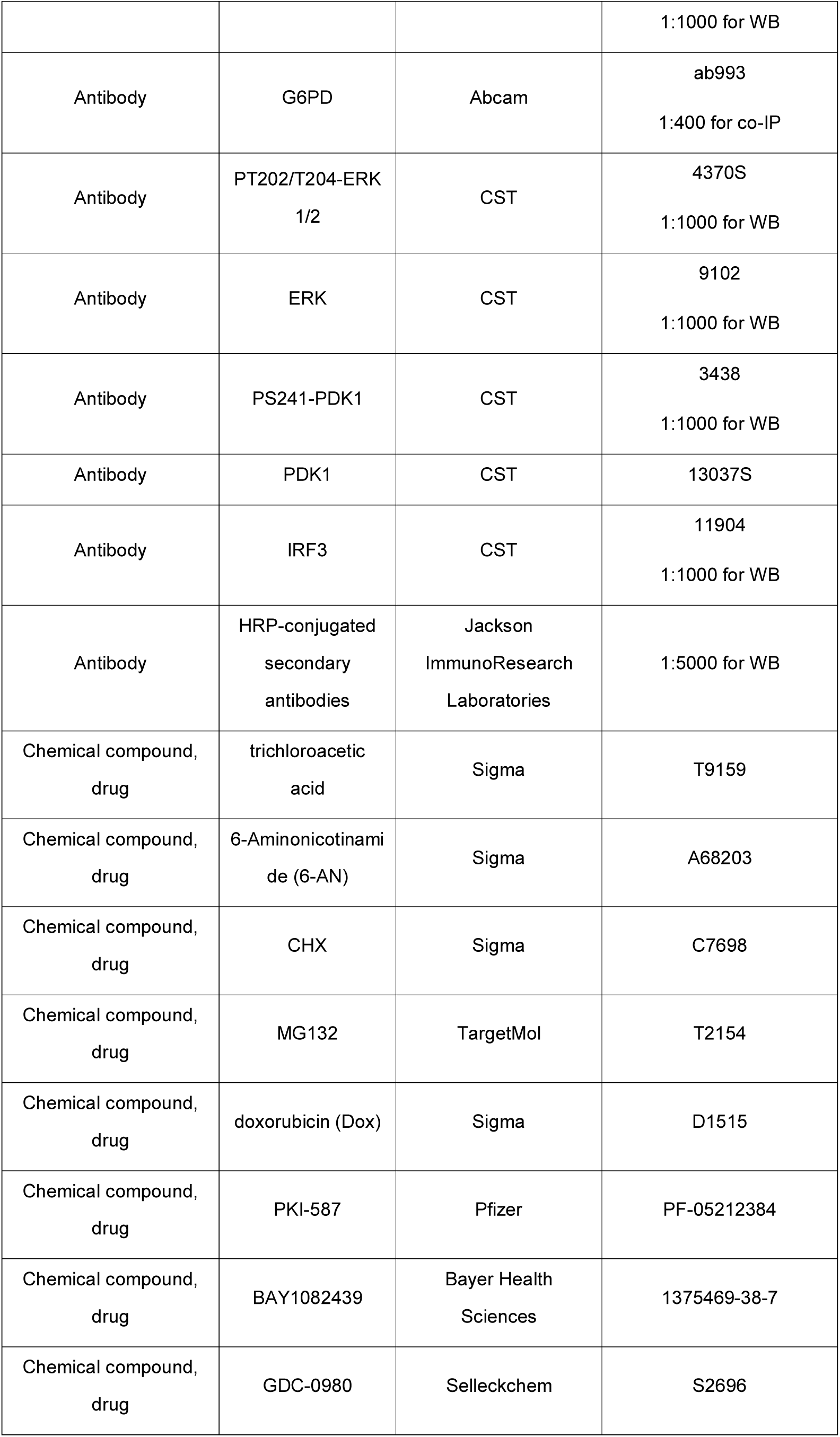
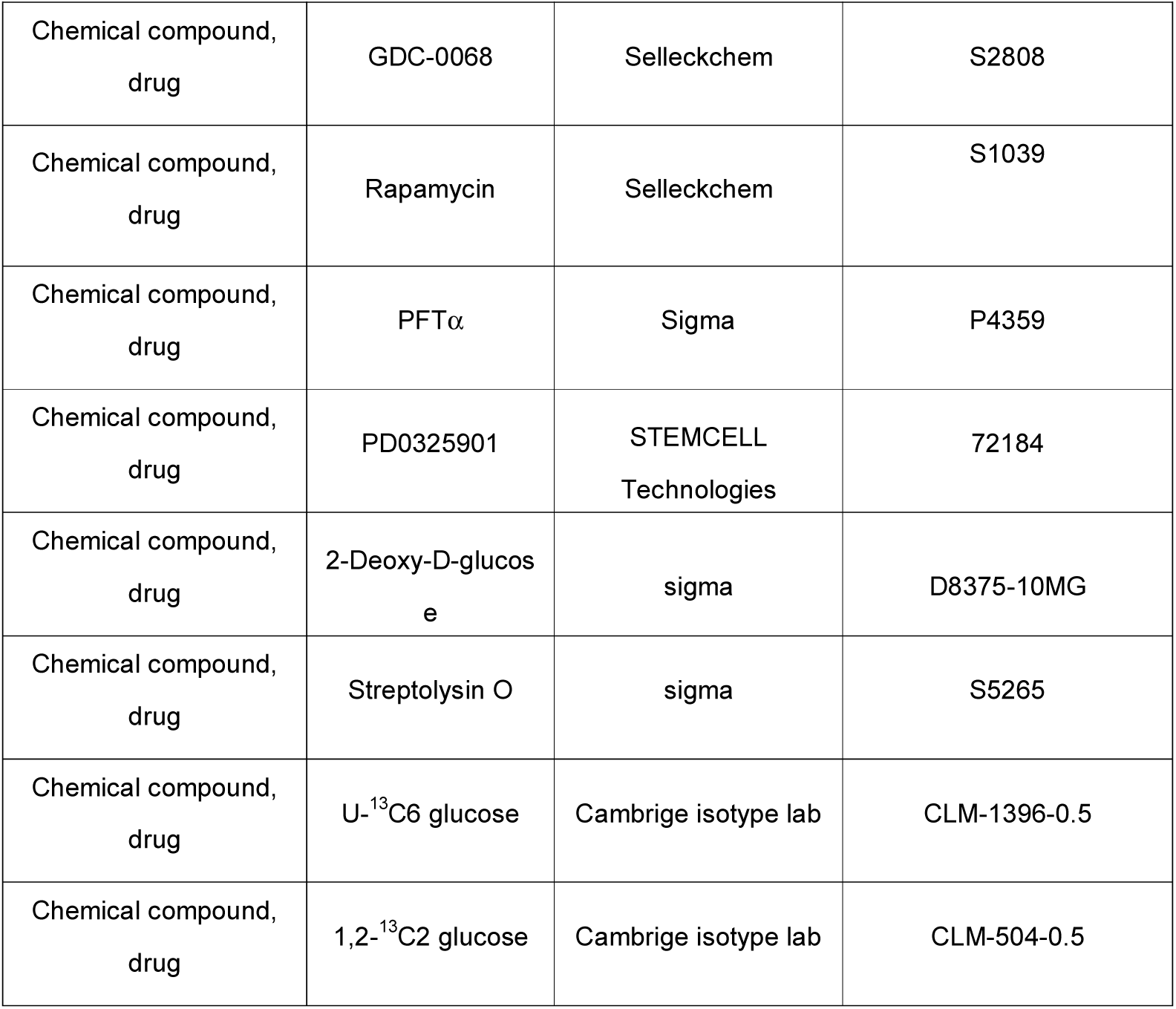

| Reagent type<br>or resource | Designation | Identifiers | Additional information |
| --- | --- | --- | --- |
| Antibody | GLUT1 | Cell Signaling<br>Technology | 12939<br>1:1000 for WB |
| Antibody | PTEN | Cell Signaling<br>Technology | 9188<br>1:1000 for WB |
| Antibody | PKM2 | Cell Signaling<br>Technology | 4053<br>1:1000 for WB |
| Antibody | Phospho-PKM2<br>(Tyr105) | Cell Signaling<br>Technology | 3827<br>1:1000 for WB |
| Antibody | AKT | Cell Signaling<br>Technology | 9272<br>1:1000 for WB |
| Antibody | Phospho-AKT<br>(Ser473) | Cell Signaling<br>Technology | 4060<br>1:1000 for WB<br>1: 800 for IHC |
| Antibody | Phospho-AKT<br>(Thr308) | Cell Signaling<br>Technology | 13038<br>1:1000 for WB |
| Antibody | S6 | Cell Signaling | 2217 |
|  |  | Technology | 1:1000 for WB |
| Antibody | Phospho-S6<br>(Ser240/244) | Cell Signaling<br>Technology | 5364<br>1:1000 for WB<br>1: for 1000 IHC |
| Antibody | GST | Cell Signaling<br>Technology | 2624<br>1:1000 for WB |
| Antibody | Histone H3 | Cell Signaling<br>Technology | 4499<br>1:1000 for WB |
| Antibody | Phospho-Histone<br>H3 (Ser10) | Cell Signaling<br>Technology | 53348<br>1:1000 for WB |
| Antibody | CASPASE 3 | Cell Signaling<br>Technology | 9665<br>1:1000 for WB |
| Antibody | Cleaved<br>CASPASE 3 | Cell Signaling<br>Technology | 9664<br>1:1000 for WB |
| Antibody | TRIM21 | Santa cruz | sc-25351<br>1:100 for co-IP, 1:1000<br>for WB |
| Antibody | ACTB | Santa cruz | sc-1616<br>1:2000 for WB |
| Antibody | G6PD | Sigma | HPA000834<br>1:1000 for WB<br>1:1000 for IHC |
| Antibody | Ki67 | Abcam | ab15580<br>1:5000 for IHC |
| Antibody | FLAG M2 | Sigma | F3165<br>1:5000 for WB |
| Antibody | HA | Sigma | H6908<br>1:5000 for WB |
| Antibody | PHLDA3 | Abcam | ab81464 |
|  |  |  | 1:1000 for WB |
| Antibody | G6PD | Abcam | ab993<br>1:400 for co-IP |
| Antibody | PT202/T204-ERK<br>1/2 | CST | 4370S<br>1:1000 for WB |
| Antibody | ERK | CST | 9102<br>1:1000 for WB |
| Antibody | PS241-PDK1 | CST | 3438<br>1:1000 for WB |
| Antibody | PDK1 | CST | 13037S |
| Antibody | IRF3 | CST | 11904<br>1:1000 for WB |
| Antibody | HRP-conjugated<br>secondary<br>antibodies | Jackson<br>ImmunoResearch<br>Laboratories | 1:5000 for WB |
| Chemical compound,<br>drug | trichloroacetic<br>acid | Sigma | T9159 |
| Chemical compound,<br>drug | 6-Aminonicotinami<br>de (6-AN) | Sigma | A68203 |
| Chemical compound,<br>drug | CHX | Sigma | C7698 |
| Chemical compound,<br>drug | MG132 | TargetMol | T2154 |
| Chemical compound,<br>drug | doxorubicin (Dox) | Sigma | D1515 |
| Chemical compound,<br>drug | PKI-587 | Pfizer | PF-05212384 |
| Chemical compound,<br>drug | BAY1082439 | Bayer Health<br>Sciences | 1375469-38-7 |
| Chemical compound,<br>drug | GDC-0980 | Selleckchem | S2696 |
| Chemical compound,<br>drug | GDC-0068 | Selleckchem | S2808 |
| Chemical compound,<br>drug | Rapamycin | Selleckchem | S1039 |
| Chemical compound,<br>drug | PFTα | Sigma | P4359 |
| Chemical compound,<br>drug | PD0325901 | STEMCELL<br>Technologies | 72184 |
| Chemical compound,<br>drug | 2-Deoxy-D-glucose | sigma | D8375-10MG |
| Chemical compound,<br>drug | Streptolysin O | sigma | S5265 |
| Chemical compound,<br>drug | U-<sup>13</sup>C6 glucose | Cambridge isotope lab | CLM-1396-0.5 |
| Chemical compound,<br>drug | 1,2-<sup>13</sup>C2 glucose | Cambridge isotope lab | CLM-504-0.5 |

### Plasmids and Transfection

The expression vectors of 3×Flag-G6PD, Flag-TRIM21, GST-TRIM21, and TRIM21 mutants were kindly provided by Prof. Peng Jiang (Tsinghua University). The HA-ubiquitin expression vector was kindly provided by Prof. Xin Ye (Institute of Microbiology, CAS). Flag-PHLDA3 vectors were constructed by cloning PHLDA3 cDNA into the pFLAG-CMV-2 vectors (Sigma). Flag-G6PD WT, Flag-G6PD K8R vectors were constructed by cloning G6PD WT or mutant (K8R) cDNA into the pFLAG-CMV-2 vectors. Myc-PTEN was made by cloning PTEN cDNA into the pCMV myc vector (Clontech). The expression plasmid of AKT WT was kindly provided by Prof. Peng Jiang. The expression plasmids of AKT mutants were generated from AKT WT plasmid. PHLDA3 promoter were cloned into pGl3.1 plasmid to drive luciferase reporter assay. PC3 or HEK293T cells were transfected with constructs for 36-48 h using Lipofectamine 2000 (Invitrogen).

### ECAR Assay

WT and *Pten* mutant mES cells were seeded in 24-well plates at a density of 2×10^4^ cells/well and cultured under standard ES culture conditions for 12 hours. Cells were then cultured in the glucose-depleted media for 1 hour and the extracellular acidification rate (ECAR) was measured after addition of glucose (80 mM), oligomycin (2 mM), and 2-DG (50 mM) using a Seahorse XF24 instrument (Seahorse Bioscience). The Seahorse XF Glycolysis Stress Test Kit was purchased from Agilent Technologies.

### RNA Interference

PC3 cells were transfected with gene-specific small interfering RNAs (siRNA) or a control siRNA (50 nM) for 72 h using Lipofectamine 2000. G6PD siRNAs were purchased from Invitrogen (Catalogue No. HSS103891). PHLDA3, and TRIM21 siRNAs were purchased from Ribobio:

siPHLDA3: GGCCCAAGGAGCTCAGCTT

siTRIM21-1^#^: GGACAATTTGGTTGTGGAA

siTRIM21-2^#^: GGAATGCATCTCTCAGGTT

### Metabolite Extraction and Mass Spectrometry-based Metabolomics Analysis

Analyses of glycolysis/PPP tracing were performed using standard procedures^19^. Briefly, cells were cultured in standard medium and then changed to medium containing 25 mM U-^13^C or 1, 2-^13^C_2_ glucose for 12 hours. Metabolites were extracted with cold 0.18 M aqueous trichloroacetic acid. Insoluble materials in lysates were centrifuged at 12,000 g for 15 min, and the resulting supernatants were freeze-dried. Samples were resuspended in 30 µl of ultrapure water for liquid chromatography-high resolution mass spectrometry^44^. The concentrations of extracted metabolites were normalized to the protein concentration measured using the BCA assay. Compounds were separated on an XBridge Amide column (100 × 4.6 mm, 3.5 μm; Waters, Milford, MA, USA). Buffer A comprised 20 mM ammonium acetate (pH 9.4) in water and buffer B was acetonitrile (LC-MS grade). All standard metabolites were purchased from Sigma-Aldrich. The retention time of ATP and GTP was very close and the concentration of GTP was close to the detection limit. Hence, we detected GDP instead of GTP.

### Co-immunoprecipitation Assays

Cells were harvested and lysed in lysis buffer at 4°C for 20 min. Cell lysates were incubated with Protein A-agarose at 4°C for 2 h followed by the addition of antibodies overnight or Flag antibody-conjugated agarose beads for 4 h at 4°C. The beads were then washed with lysis buffer. The immunoprecipitants were eluted from the beads with 2×SDS loading buffer, separated on 10% SDS-PAGE gels, and immunoblotted with the indicated antibodies.

### Enzymatic Activity Measurements

The pyruvate kinase activity assay was performed using a pyruvate kinase activity assay kit (Sigma, catalog number MAK072) according to the manufacturer’s protocol. The G6PD enzyme activity was determined as previously described ^45^.

### Luciferase assay

PC3 cells were transfected with luciferase reporter plasmids or control plasmid, pGL3.1, for 24 h. The Steady-Glo® Luciferase Assay System (Promega) was used to quantify luminescence from transfected cells.

### Colony Formation Assay

WT or *Pten* null mES cells were seeded in 3.5 cm dishes at a density of 2,000 cells/well. After 24 h, cells were treated with different drugs. After 4 days of drug treatment, cells were fixed with methanol for 30 min, stained with trypan blue for 30 min at room temperature, and then the colonies with a diameter greater than 100 mm were counted.

### Cell Viability Assay

*TRIM21* WT and KO cells or *PHLDA3* WT and KO cells were seeded in 96-well plates. After 24 h, *PHLDA3* WT and KO cells were treated with 6-AN, R5P (2 mM), or uridine (2 mM) for 4 days. The CCK-8 reagent (YEASEN) was added to each well and incubated for 1 h according to the manufacturer’s protocol. The mean absorbance at 450 nm was measured.

### Permeabilization with SLO

G6P treatment was accompanied by SLO permeabilization. PC3 cells were permeabilized using a procedure similar to a previously described method ^46^. Briefly, PC3 cells were washed three times with IC buffer (140 mM potassium glutamate, 20 mM Hepes, 5 mM MgC1_2_, 5 mM EGTA, and 5 mM NaCl, pH 7.2) and incubated in IC buffer containing 0.5 IU/ml SLO and 2 mM G6P for 10 min at 37°C to permeabilization. Then cells were changed with normal medium for the indicated times.

### Cell Apoptosis Assay

The collected cells were subjected to apoptosis detection using flow cytometry on a BD Accuri C6 flow cytometer (BD Biosciences, San Jose, CA, USA) using the Annexin V and PI apoptosis detection kit (MultiSciences), according to the manufacturer’s protocol.

### Cell Cycle Assay

Cells were harvested and fixed with ethanol overnight at -20°C before staining with propidium iodide. Flow cytometry was performed with a FACScan instrument.

### Xenograft tumor formation assays

2.5×10^6^ *TRIM21* WT or isogenic *TRIM21* knockout cells were mixed 1:1 in LDEV-free matrigel (Corning, SLS 354234) and injected subcutaneously into bilateral flank of male CAnN.Cg-Foxn1^nu^/CrlVr mice Xenograft tumors were measured twice every six days and tumor volume was calculated using the following calculation: volume (mm^3^) = (length x width x width) / 2.

For 6-AN treatment, equal number of *PHLDA3* WT or isogenic *PHLDA3* knockout cells (2.5×10^6^) were mixed 1:1 in matrigel and implanted onto bilateral flank of nude mice. The xenograft growth was monitored daily. When becoming palpable, mice were treated with 1 mg/kg 6-AN once daily IP for 30 days. 6-AN was freshly prepared in 1% DMSO in PBS. Xenograft tumors were measured every six days with a caliper and tumor volumes and weight were calculated as above.

### Reverse Transcription and PCR Analysis

Total RNAs and cDNAs were prepared using TRIzol (Invitrogen) and qPCR (Vazyme R223-01). PCR was performed according to the manufacturer’s instructions (Vazyme Q121-03).

The following primer pairs were used for quantitative real-time PCR:

Human GAPDH (forward): 5’-GGGGAGCCAAAAGGGTCATCATCT-3’

Human GAPDH (reverse): 5’-GAGGGGCCATCCACAGTCTTCT-3’

Mus Actb (forward): 5’-GGGGAGCCAAAAGGGTCATCATCT-3’

Mus Actb (reverse): 5’-GGGGAGCCAAAAGGGTCATCATCT-3’

Human PHLDA3 (forward): 5’-CCGTGGAGTGCGTGGAGAGC-3’

Human PHLDA3 (reverse): 5’-CTAGGGTGATCTGGGCGTTCC-3’

Mus Phlda3 (forward): 5’-CCGTGGAGTGCGTAGAGAG-3’

Mus Phlda3 (reverse): 5’-TCTGGATGGCCTGTTGATTCT-3’

Human TRIM21 (forward): 5’-TCAGCAGCACGCTTGACAAT-3’

Human TRIM21 (reverse): 5’-GGCCACACTCGATGCTCAC-3’

Mus Trim21 (forward): 5’-TGGTGGAGCCTATGAGTATCG-3’

Mus Trim21 (reverse): 5’-GGCACTCGGGACATGAACTG-3’

Mus G6pd (forward): 5’-CACAGTGGACGACATCCGAAA-3’

Mus G6pd (reverse): 5’-GCAGGGCATTCATGTGGCT-3’

### Western Blot

Cells were harvested, lysed in RIPA buffer (1% SDS) containing protease inhibitors (Roche Applied Science), and quantified using a BCA Protein Assay Kit (Pierce). Cell extracts were separated on 10% SDS-PAGE gels, transferred to PVDF membranes (Millipore), blocked with Tris-buffered saline/Tween 20 (TBST) containing 5% skim milk for 1 h at room temperature, and then probed with the indicated primary antibody at 4°C overnight. After three washes, membranes were incubated with a 1:5,000 dilution of HRP-conjugated secondary antibodies for 1 h at room temperature. The immunoblots were detected using ECL (Thermo) according to the manufacturer’s protocol.

### Statistical Analysis

All quantitative data are presented as the means ± SD of at least three independent experiments. Statistical significance was determined by Student’s *t* -test and expressed as a P value.

Normalized and log2 transformed gene expression data (RNA sequencing) of TCGA Pan-Cancer were downloaded from Xena data hub (https://xenabrowser.net/hub/). The pathway activity score was calculated by PLAGE^47^. The correlation was tested by Spearman’s rank correlation test.

